# Late-Stage Large Extracellular Vesicles Reprogram CHO Cell Metabolism in a Glutamine-Dependent Mode and Promote Antibody-Productivity to Cell-Growth Tradeoff

**DOI:** 10.64898/2026.06.28.735077

**Authors:** Hong B. Nguyen, Nikola G. Malinov, Alekhya Puttagunta, Kelvin H. Lee, Eleftherios T. Papoutsakis

## Abstract

Extracellular vesicles (EVs) are mediators of intercellular communication, yet their impact on Chinese Hamster Ovary (CHO) cell physiology and bioprocess performance remains poorly understood. Here, we investigated whether small EVs (sEVs) and large EVs (LgEVs) that accumulate during fed-batch and perfusion cultures modulate CHO cell growth, metabolism, apoptosis, and monoclonal antibody (mAb) production. EVs isolated from early- and late-stage cultures were added to fresh CHO cultures grown with or without glutamine supplementation. Only LgEVs had a significant impact. Late-stage LgEVs markedly altered CHO-cell behavior, reducing cell proliferation, increasing apoptosis under glutamine-limited conditions, and substantially enhancing mAb productivity in a dose-dependent manner. Glutamine supplementation largely alleviated the growth-inhibitory and pro-apoptotic effects of LgEVs while preserving their positive impact on productivity, suggesting that glutamine decouples EV-mediated stress from productivity enhancement. Metabolic analyses revealed increased glucose consumption, a glutamine-dependent shift between glycine and alanine overflow metabolism, and remodeling of amino-acid utilization. Metabolic flux analysis further demonstrated enhanced glycolytic overflow and increased reliance on amino acid-supported anaplerosis. Conversely, selective removal of LgEVs from perfusion medium significantly improved cell expansion without reducing antibody production, supporting an inhibitory role for late-stage LgEVs. These LgEVs were enriched in let-7 family miRNAs and miR-21, consistent with RNAseq analyses demonstrating stress-associated enrichment of these miRNAs in CHO EVs and with functional studies showing that let-7a and miR-21reduce CHO-cell growth. Together, these observations suggest that selective miRNA loading contributes to the growth, metabolic, and productivity phenotypes elicited by late-stage LgEVs. Our findings identify LgEVs as endogenous regulators of CHO-cell physiology and potential targets for optimizing high-density fed-batch and perfusion biomanufacturing processes.

**Highlights:**

- Endogenous late-stage Large Extracellular Vesicles (LgEVs) reduce CHO cell growth but boost specific mAb productivity.
- Glutamine supplementation rescues LgEV-mediated growth inhibition and apoptosis.
- Metabolic Flux Analysis (MFA) based on the dynamic behavior of amino acid and other metabolite and substrate concentrations reveals the pyruvate node as a metabolic bottleneck and the associated lactate overflow metabolism as resulting from LgEV exposure.
- Stress-associated let-7 and miR-21 microRNAs are highly enriched on a per-EV basis in late-stage LgEVs.
- Selective removal of LgEVs improves perfusion cell growth without impacting antibody titer.

## I. Introduction

Chinese Hamster Ovary (CHO) cells are used to produce most biotherapeutic proteins – including monoclonal antibodies (mAbs) – a fast-growing market expected to reach and exceed one trillion dollars by 2032. Despite the enormous progress achieved since the first approved product processes in the late 1980s, the drive to understand and improve CHO culture performance continues unabated, now with increased emphasis on continuous perfusion biomanufacturing. Since their first introduction in 1958 as “10 months with no diminution in growth rate or change in cellular or colonial morphology” (Puck et al., 1958)CHO cells have been the reliable model cell line in the industry for several reasons, including robust and reliable growth, and their ability to perform human-like post-translational modifications (Puck et al., 1958; Szkodny & Lee, 2022). To meet the demand for affordable therapeutics, further improvements in mAb production depend not only on refining process conditions but also on deepening our understanding of how cells adapt over time and which culture components can influence overall process outcomes. While metabolic inhibitors like ammonia, lactate, and formate are well-studied, they represent only a fraction of the ‘secretome’ (McCartney et al., 2019; Mulukutla et al., 2017). Beyond these small molecules lies a complex layer of intercellular communication mediated by extracellular vehicles (EVs), which are constantly generated and accumulate in CHO cultures (Belliveau & Papoutsakis, 2022, 2023; Busch et al., 2022; Keysberg et al., 2021). Notable is, the finding that EVs dynamically mediate massive exchange of cellular material (including RNA and proteins) between cells under normal culture conditions (Belliveau & Papoutsakis, 2022), and that the EV cargo, notably miRNA cargo, changes dynamically in response to culture conditions, including metabolic stresses (Belliveau & Papoutsakis, 2023).

EVs are lipid bilayer particles that carry proteins, small RNA [miRNA, lncRNA, piRNA], and sometimes DNA. EVs are typically categorized by their biogenesis—exosomes (50 to 150 nm) via the endosomal pathway and microparticles (150 to 1000 nm) via outward membrane budding—but their functional impact is driven by the proteins and RNAs they shuttle between cells (Bazzan et al., 2021; Van Niel et al., 2018). Due to their considerable overlap in sizes and isolation via differential ultracentrifugation, we categorized them as small EVs (sEVs: enriched in exosomes) and large EVs (LgEVs: enriched in microparticles) per the International Society of Extracellular Vesicles (ISEV) guidelines (Welsh et al., 2024). The functional significance of EVs is increasingly recognized in the context of physiology and disease, with roles in immune regulation, cancer metastasis, wound healing, and platelet biogenesis (Escobar et al., 2020; Jiang et al., 2017; O. P. B. Wiklander et al., 2019; Oscar P. B. Wiklander et al., 2019).

Despite a growing body of literature, EVs remain largely overlooked in bioprocessing, even though they are produced in large quantities and continuously exchanged between cells throughout culture (Belliveau & Papoutsakis, 2022). EV RNA (and notably miRNAs) and protein contents vary significantly with culture stage or stress conditions in CHO-cell cultures (Belliveau & Papoutsakis, 2023; Busch et al., 2022; Keysberg et al., 2021; Kumar & Banerjee, 2016). Yet, little is known about how EVs influence culture performance metrics—such as nutrient consumption, cell proliferation, and mAb productivity—under industrially relevant fed-batch or perfusion conditions. Investigating EVs in CHO systems, therefore, offers an opportunity to uncover new layers of cellular communication that directly impact culture performance and therapeutic protein yield. We hypothesized that EVs carry growth-promoting components during the exponential phase and growth-inhibiting components at later stages. Given the massive exchange of cellular material they mediate, EVs from different culture stages and conditions are likely to affect cell metabolism and phenotype (Belliveau & Papoutsakis, 2022; Thompson & Papoutsakis, 2023). Here, we investigated the biological activity of CHO-derived EVs collected at distinct culture stages: exponential growth, stationary phase, and at the onset of filter fouling in perfusion cultures. We examined whether and how these EVs alter CHO cell metabolism by transferring their cargo to recipient cells. Using MFA, we focused on the impact of core metabolic reactions that support cell growth and energy production, as well as how they might affect cell growth, death, and antibody productivity. Although EV-mediated metabolic reprogramming has been increasingly recognized, particularly in tumor microenvironment studies, few investigations have quantified detailed EV-induced metabolic changes, and even fewer have done so at the level of intracellular metabolic fluxes using Metabolic Flux Analysis (MFA). Most published studies have relied on measurements of glucose and glutamine consumption, lactate production, oxygen consumption, extracellular acidification, metabolomics, or expression of metabolic enzymes and transporters rather than formal metabolic flux analysis (Fridman et al., 2022; Jiang et al., 2023).

## II. Materials and Methods

### 2.1 CHO cell cultures: batch, fed-batch, and perfusion

The CHO-K1 cell line (Clone A11 from the Vaccine Research Center at the National Institutes of Health), stably expressing anti-HIV IgG 1 (VRC01), was used in this study. CHO VRC01 cells were thawed and passaged in 50-mL spin tubes with 20 mL working volume. HyClone ActiPro media (Cytiva) was used as a basal medium throughout the study. Since basal media are formulated without L-Gln or Insulin, 6 mM Gln was supplemented at the start of cultures or at passage. Seeding density for batch, fed-batch, and perfusion cultures was 0.4 million cells/mL. Batch and fed-batch cultures were carried out in 125 mL and 500 mL shake flasks (Corning, vented cap), with initial working volumes of 25 mL and 80 mL, respectively, to ensure sufficient oxygen transfer. The incubator was set at 37°C, 5% CO2, 20% O2, 85% RH, with 190 RPM on a 25-mm diameter orbital shaker (Infors HT Multitrons). In fed-batch cultures, bolus feeding protocols were employed, in which HyClone Cell Boost 7a and 7b (Cytiva), rich in essential amino acids, were added daily from day 3 onward at 3% v/v and 0.3% v/v, respectively. From day 5 onward, supplementary glucose (450 g/L) was added daily after feeding Cell Boost 7a and 7b to target a glucose concentration of 9 g/L.

Continuous perfusion cultures were recently described (Malinov et al., 2026). In brief, perfusion cultures were performed in a 1-L Eppendorf BioFlo® 320 bioreactor. Cell retention was achieved via tangential flow filtration (TFF) using a centrifugal pump (PuraLev® i30MU) and hollow fiber modules (Repligen): either a 0.65 μm mPES module (520 cm²) or a 0.2 μm PES module (1000 cm²), both 20 cm in length. Culture began in batch mode and transitioned to perfusion on day 3 at one vessel volume per day (vvd^-1^). The perfusion rate was controlled by the harvest line pump and gravimetric feeding to maintain a constant working volume. Steady state was reached at a viable cell density (VCD) setpoint of 40 million cells/mL, maintained via a daily intermittent cell bleed. The temperature was controlled at 37°C. The pH was adjusted to 7.00 ± 0.05 via CO₂ sparging. Dissolved oxygen (DO) was maintained at 40% of air saturation using three-gas control (air, CO₂, O₂), with sparging rates from 0.02–0.40 SLPM and a constant 0.05 SLPM air overlay. Agitation began at 90 rpm with a pitch-blade impeller and increased as needed to meet oxygen demand. Antifoam C emulsion solution (5%) (Sigma Aldrich) was added as required. Daily samples were collected for offline analysis, including pre- and post-bleed during steady state. Perfusion EVs were harvested via daily cell bleeds on Day 8, and 11 of the same runs as the previous publication (Malinov et al., 2026).

### 2.2 EV isolation and counting

EVs were harvested from cultures with viability above 90% via previously described and characterized differential ultracentrifugation of CHO-VRC01 EVs (Belliveau & Papoutsakis, 2022, 2023). Isolated CHO EVs were resuspended in fresh basal culture media for immediate use or aliquoted for −80°C storage. CHO LgEVs were counted using a CYTOFLEX S flow cytometer in Violet Side Scatter mode (VSSC) and analyzed with CytExpert software (Beckman Coulter). Particles with diameters between 200 nm and 1000 nm can be gated using fluorescent calibration beads (Invitrogen, Cat # F13839). Before counting, the CYTOFLEX S underwent overnight deep cleaning, followed by daily cleaning, and a 0.2 μm-filtered PBS flush to ensure a clean baseline. VSSC was set up according to the manufacturer’s protocol on the CYTOFLEX S. CHO sEVs were counted using Nanoparticle Tracking Analysis (Nanosight NS3000, Malvern Panalytical) at a 500–1000-fold dilution in freshly filtered PBS (0.2 μm). Data acquisition and analysis were performed using NTA software version 3.3, with the camera level set to 13-14, a detection threshold of 5, and triplicate measurements at 30-second intervals. Particle concentrations were recorded as the average of the three replicates.

### 2.3 EVs addition to fresh cultures

For EV-supplemented cultures, using a 24-well deep-well plate (Axygen), CHO cells were inoculated at 0.4 million viable cells per mL into each well, with a working volume of 2.5 mL in a total volume of 10 mL under the same incubator settings as above, with a shake speed of 200 RPM on a 25-mm throw-diameter orbital shaker. EVs resuspended in fresh medium were added at the desired dosing concentration. Based on LgEV and sEV concentrations typically found in fed-batch and perfusion cultures, low, mid, and high doses of EVs were chosen to be 2×10^7^, 4×10^7^, and 1×10^8^ LgEVs per mL culture and 4×10^9^- 4×10^10^sEVs per mL culture, respectively. For some experiments, 6 mM Gln was added during inoculation (**Gln^+^**). Viable cell density and viability were assessed using trypan blue exclusion on the Vicell-XR.

### 2.4 Flow cytometry for cell apoptosis

Cell apoptosis was determined by flow cytometry using two detection methods when applicable, Annexin V and 7-aminoactinomycin D (7AAD) (BioLegend), and Caspase 3/7 (CellEvent™ Caspase-3/7 Green Detection Reagent). Activation of Caspase 3 inactivates the function of flipase ATP11, which in turn exposes phosphatidylserine (PS) on the outer leaflet of the cell, marking apoptosis (Nossing & Ryan, 2023). Briefly, cells were washed twice with cold PBS and resuspended in Annexin V Binding Buffer. 2 µL of FITC-Annexin V and 2 µL of 7AAD were added to 100 µL of the cell suspension, followed by incubation for 15 min. Caspase 3/7 reagent is a cell-permeant, fluorogenic substrate consisting of an amino acid peptide conjugated to a nucleic acid-binding dye. When caspases 3 and 7 are activated in apoptotic cells, they cleave the nucleic acid-binding dye. For caspase activation detection, 200 μL of cell suspension was incubated with 1 μL of detection reagent at 37°C for 45 min. No wash or resuspension is needed. Cells were analyzed using the CYTOFLEX S flow cytometer at a constant flow rate. Cells were gated to exclude doublets using FSC-H vs FSC-W, and the percent specific cell population was analyzed with at least 20,000-40,000 total cell events per sample.

### 2.5 Measurement of substrate, metabolite, and IgG concentrations

Glucose, lactate, and ammonia measurements were performed on a YSI 2950 bioanalyzer. Concentrations of 18 amino acids were measured using an AdvancedBio Amino Acid analysis column (Agilent, 655950-802) with a UHPLC Guard column (Agilent, 820750-931) on an Agilent HPLC 1260 Infinity II as described previously (Reddy et al., 2025). Antibody titers were measured using a POROS Protein A column (Cat# 2-1001, 2.3cm column) as described (Reddy et al., 2025). HPLC-grade IgG from a human serum standard (Sigma-Aldrich, Cat #I2511-10mg) was used for the calibration curve. Specific amino acid and metabolite consumption and production rates were calculated by dividing the residual concentration differences between the days by the integral viable cell density (IVCD) of the cultures.

IVCD is the area under the growth curve and is approximated using the trapezoid method:

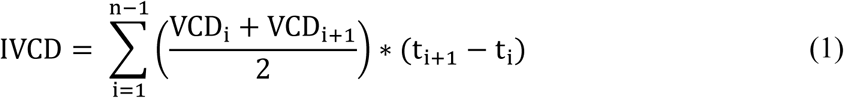

Specific metabolite consumption/production rate (mg/10⁶cells/day) and productivity (pg/cell/day):

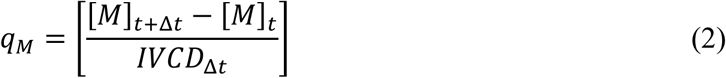

### 2.6 Media fractionation and satellite cultures

Spent media fractionation was performed by two key methods, ultracentrifugation and ultrafiltration. Satellite cultures are subcultures derived from ongoing CHO perfusion or fed-batch cultures, in which the same perfusion cells are resuspended in different media fractions. The schematic of the experimental setup is shown in Figure S1. On the day of media fractionation, cells were first removed by centrifugation at 180 × g for 5 min. Before media processing, HyClone Cell Boost 7a and 7b were added at 3% v/v and 0.3% v/v, respectively, to ensure adequate nutrient availability. Apoptotic bodies from spent media were removed by centrifugation at 2,000× g for 15 min. The supernatant from the 2,000× g spin was subsequently ultracentrifuged at 25,000× g for 30 min to remove LgEVs. The supernatant was then subjected to 100,000× g ultracentrifugation for 90 min at 4°C to remove sEVs. Supernatants from either 2,000× g spin were loaded into a 50 mL Amicon ultra centrifugal filter (100 kDa MWCO, Millipore) and centrifuged at 3,000× g for 1.5 hours at 37°C, until 1 mL of the retentate remained. Filtrate from a 100-kDa filter was added to a 50 mL Amicon ultra centrifugal filter 5-kDa MWCO (Millipore) and centrifuged at 3,000× g for 1.5 hours. Glucose and lactate were measured in each fraction, revealing no significant differences in core metabolite levels across fractions. Conditioned cells from each day were seeded at a target of 5 million cells per mL in the corresponding media fractions from the same day. Cells were pelleted at 180× g for 5 minutes before being resuspended in the pre-warmed fractionated media. Cell counts were performed daily on Vi-cell XR. HyClone Cell Boost 7a and 7b were fed at 5% v/v and 0.5% v/v, respectively, to ensure nutrient availability.

### 2.7 Extraction of RNA and reverse transcription quantitative PCR (RT-qPCR) of selected miRNAs

Quantification and RT-qPCR of specific miRNAs were performed as described (Kao et al., 2022; Thompson & Papoutsakis, 2023). Briefly, miRNA was isolated using the miRNeasy micro kit (QIAGEN, cat #217084) per the manufacturer’s protocol. One-half picomole of synthetic cel-miR-39-3p (Thermo Fisher Scientific) was added to samples during the cell or EV lysis step as a spike-in control, enabling calculation of miRNA copy number per EV. Isolated RNA concentration was measured with the Qubit RNA concentration kit (Qiagen, cat# Q32855). Then, the appropriate amount of RNA was reverse transcribed using the TaqMan MicroRNA Reverse Transcription Kit with pre-optimized specific primer/probe sets from TaqMan MicroRNA assays (Thermo Fisher Scientific, cat# 4427975: hsa-miR-92a, hsa-miR-21, hsa-let-7a, hsa-let-7b, hsa-let-7c). qPCR assays were carried out using the TaqMan Universal PCR Master Mix II with added miRNA-specific probe as above. Technical triplicates were performed for each biological replicate in qPCR analysis. The C1000 Touch ThermoCycle and CFX96 Optical Reaction Module (Bio-Rad) were used for reverse transcription and qPCR, respectively. With a known amount spiked in miR-39, assuming the purification and amplification efficiencies were the same for all miRNAs, the 2^(-ΔΔCt)^ method was used to calculate the total copy number of the specific miRNA present in the sample (Livak & Schmittgen, 2001; Thompson & Papoutsakis, 2023). The calculated miRNA copy number was normalized to the initial total number of cells or EVs used for RNA isolation.

### 2.8 Metabolic Flux Analysis

Metabolic flux analysis (MFA) was performed using a previously developed reduced metabolic-reaction model for the CHO-K1 VRC01 cell line, consisting of 42 metabolites and 68 reactions (Venkatarama Reddy et al., 2026). Metabolic flux analysis was formulated as a weighted least squares problem solved with the DAQP solver (Arnström et al., 2022) as the backend quadratic programming solver from Python’s qpsolvers library (Caron et al., 2026). The weighted sum of squared residuals between the MFA-calculated exchange fluxes, 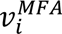, and the experimentally measured exchange fluxes, 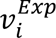, for every measured species, *_i_*, was minimized (Equation 1).

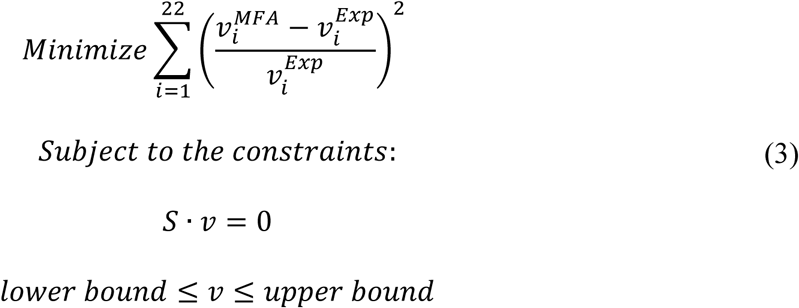

The objective function was constrained by the pseudo-steady-state assumption, where S is the stoichiometric matrix obtained from the reaction network and *_v_* is the reaction flux vector. The exchange reactions for biomass, mAb, glucose, lactate, and 18 amino acids (22 species total) were constrained to fall between 80% and 120% of their experimentally measured values. If the network was overconstrained, the upper and lower bounds were increased by 20% increments until a feasible solution was found. Ammonia was not measured for the datasets used for MFA; however, the ammonia exchange reaction was constrained to the minimum and maximum experimental values determined from prior fed-batch experiments (data not shown). Intracellular reactions were constrained to between −∞ and +∞ for reversible reactions and 0 and +∞ for irreversible reactions as described previously (Venkatarama Reddy et al., 2026).

### 2.9 Statistical analysis

Data were processed and figures generated using GraphPad (v.10.1.2). Data are presented as the mean ± standard error of the mean (SEM) of biological replicates (n ≥ 3), unless otherwise stated. Unpaired Student t-tests were performed with *p < 0.05, **p < 0.01, and ***p < 0.001.

## III. Results & Discussion

### 3.1 EVs concentration in batch, fed-batch, and perfusion cultures increases over culture time

Across all culture types, the concentrations of both sEV and LgEV subtypes increased significantly over the culture duration (Figures 1B and 1C). Their concentration increases 5–10-fold every 3 days, with LgEVs reaching peak concentrations of 5 × 10⁷ LgEVs/mL on day 8 of batch cultures and 2 × 10⁸ LgEVs/mL on day 11 of perfusion cultures. To determine whether the accumulation was due to increased cell densities, we normalized EV counts to the viable cell density (VCD). While early days production remained relatively stable (0.5 LgEVs/cell, 500 sEVs/cell), normalized levels spiked nearly eight-fold at later stages (reaching 4 LgEVs/cell and >3000 sEVs/cell) on day 8 of fed-batch and day 11 of perfusion, indicating that late-stage cells undergo a physiological shift that increases their EV secretory footprint.

**Figure 1.**
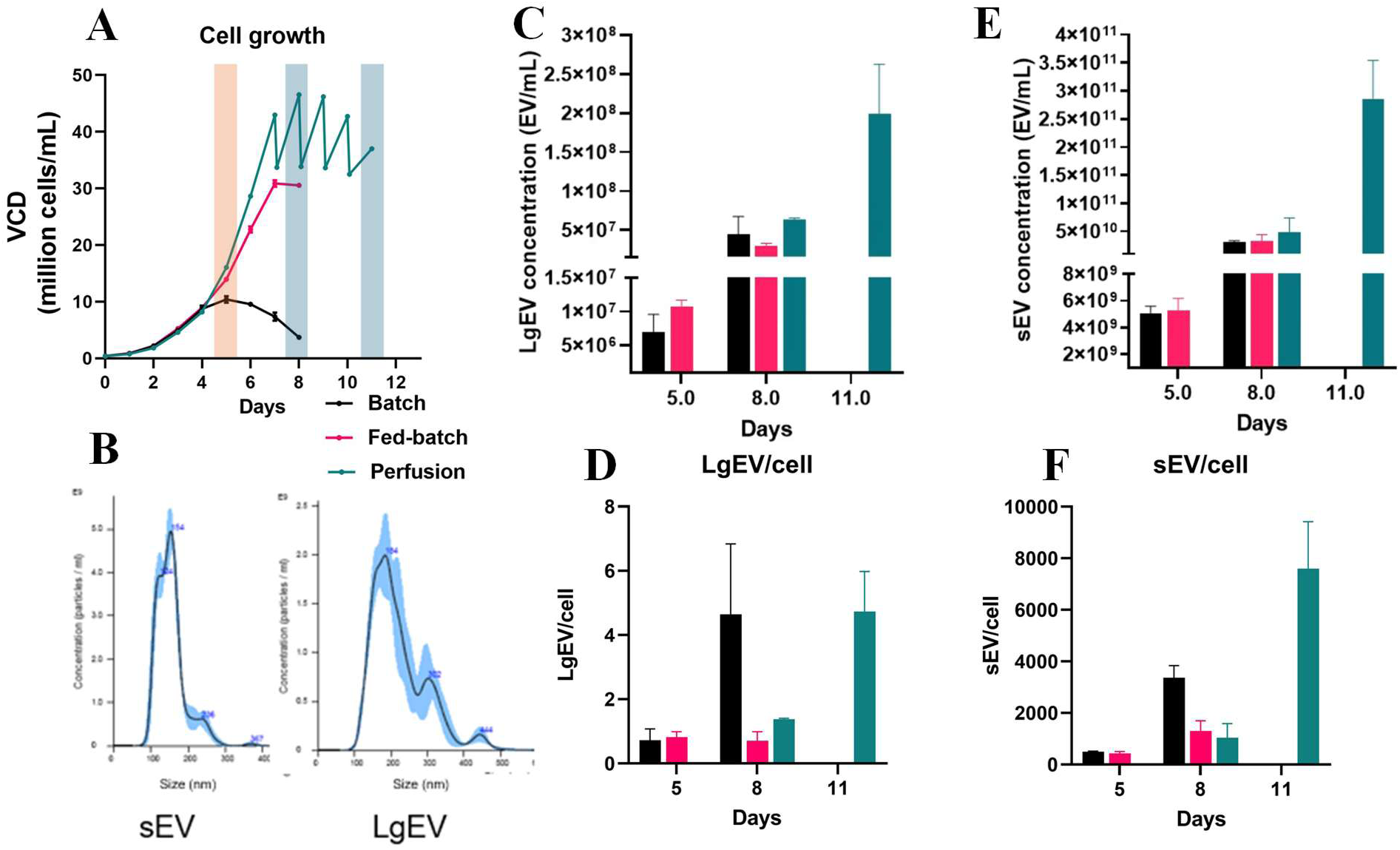
CHO cell growth and EVs counts in batch, fed-batch (in shake flasks), and perfusion cultures (in bench-top bioreactor). (A) CHO cell growth; blue bars indicate where EVs in fed-batch and perfusion cultures; light orange bar indicates the time where EVs were collected in batch and fed-batch culture. (B) NTA size characterization of sEV and LgEV of D5 fed-batch cultures. (C) LgEVs count via flow cytometry per mL of culture fluid, and (D) per viable cell. (E) sEVs count via NTA per mL of culture fluid; and (F) and per viable cell;

Error bar for batch, fed-batch cultures represent biological replicates n ≥ 3. Error bar for perfusion culture represents technical replicates of samples collected from one bioreactor run; day 8 to day 11 represented steady-state perfusion operation according to viable cell counts.

Our sEV concentrations were consistent with previously reported batch culture values (Belliveau & Papoutsakis, 2022). However, our LgEV counts were approximately three orders of magnitude lower than those reported in studies using Nanoparticle Tracking Analysis (NTA). The discrepancy is likely due to the detection limits of flow cytometry (CytoFLEX S), which only reliably capture particles >180-200 nm, whereas NTA includes smaller particles down to 50 nm (George et al., 2021). Notably, when compared to other CHO studies using flow cytometry, our LgEV production rates per cell are highly consistent (Zavec et al., 2016), validating our dosing conditions for subsequent addition experiments.

### 3.2 LgEVs from late stages of cultures inhibited cell growth but increased cell-specific productivity

To explore the biological activity of CHO-derived EVs, isolated LgEV and sEV were added to fresh cultures (“treatment”) at concentrations representative of those observed in cultures. While the addition of sEVs yielded no significant functional impact (data not shown), the addition of LgEVs from the later stages of fed-batch and perfusion cultures reduced cell growth while increasing cell-specific productivity (Figure 2). These experiments were conducted under glutamine-free (**Gln^-^**) conditions to reflect industrially relevant states where glutamine is typically depleted by day 3 of culture and is not replenished, aiming to enhance antibody production and prevent ammonia accumulation (Fomina-Yadlin et al., 2014; Hong et al., 2010; Kim et al., 2013).

**Figure 2.**
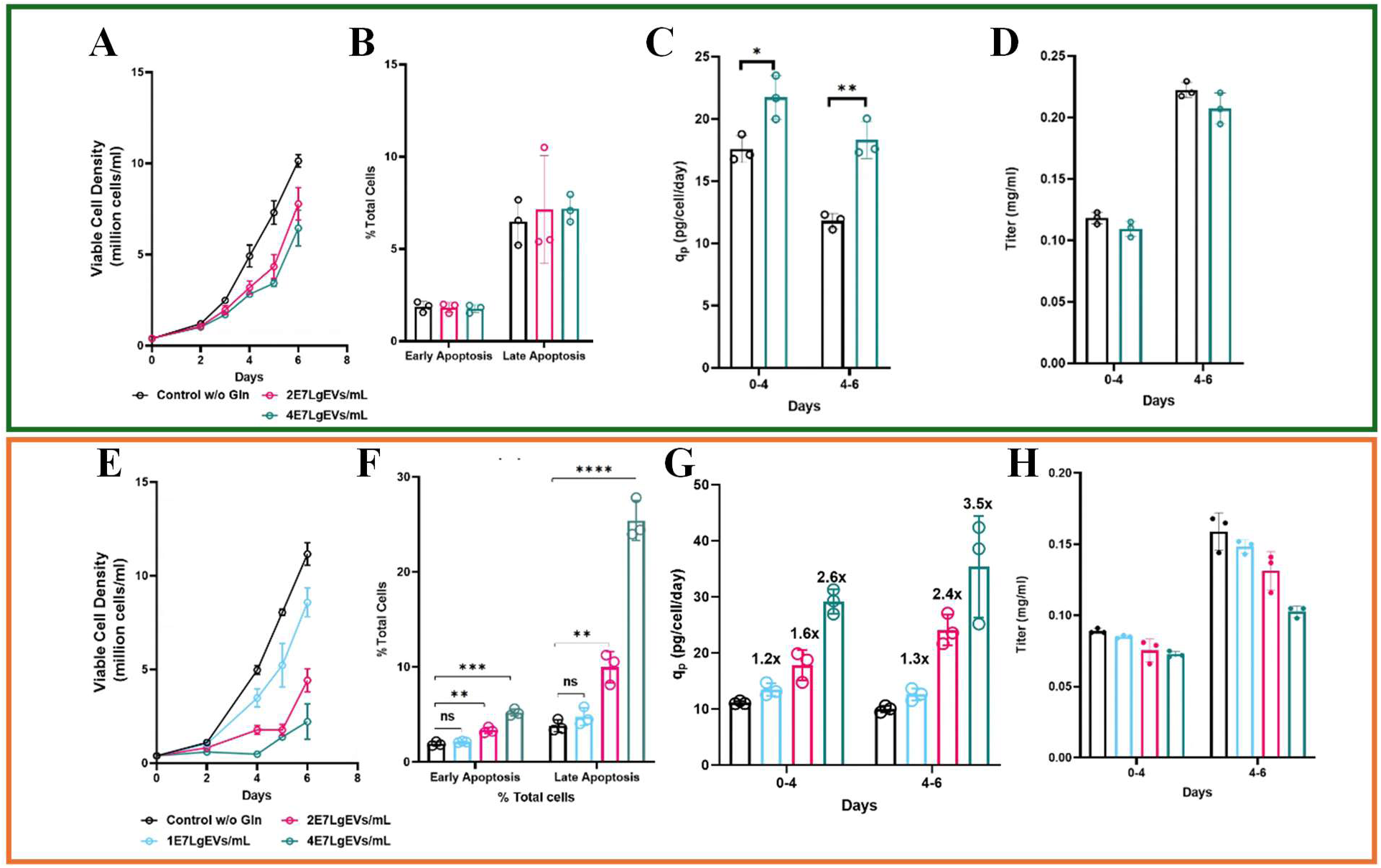
Impact of different dosages of LgEVs from D8 fed-batch cultures (panels A-D) and D11 perfusion culture (panel E-H) on cell growth, productivity, and apoptosis of fresh CHO cultures without glutamine addition (Gln^-^). (A-D): Gln^-^ cultures after adding D8 fed-batch LgEVs (2×10^7^, 4×10^7^ LgEVs/mL) or no EV addition (control). (E–H) Gln^-^ cultures after adding D11 perfusion LgEVs (1×10^7^, 2×10^7^, 4×10^7^ LgEVs/mL) or no EV addition (control). (A, F) Viable cell density (million cells/ mL) over 6 days after adding EVs at day 0. (B, F) Percentage of early and late apoptotic cells at day 4 of cultures as assessed by AnnexinV/7AAD, and by Caspase 3/7 flow cytometry assays. (C, G) Cumulative Antibody titer (mg/ml) from day 0 to day 4 and from day 4 to day 6. (D, H) Cell-specific antibody productivities (q_p_): day 0 to day 4 and day 4 to day 6 (pg/cell/day). Error bars represent standard deviation of biological replicates (n=3).

Under these conditions, Day 8 fed-batch LgEVs (4 × 10⁷ LgEVs/mL) did not affect cell viability but reduced cell growth by 35% compared to the control by day 4 post-treatment. While VCD was reduced, treated cultures showed a marked increase in cell-specific antibody productivity (q_p_), rising by 23% in the first 4 days and peaking at a 55% increase from day 4–6 (Figure 2D). The trend was even more pronounced with LgEVs from day 11 perfusion cultures, which inhibited growth and increased productivity in a dose-dependent manner. At the highest dose, 4 × 10⁷ LgEVs/mL, the maximum VCD reached only 2 million cells/mL, compared with 11.5 million cells/mL in controls. The reduction in density was closely tied to a spike in cell death, assumed to be largely to be apoptosis. Specifically, early and late apoptotic populations reached 5.6% and 25.4%, respectively, compared to 2.1% and 3.8% in controls (Figure 2F). While lower LgEV doses suppressed growth without a significant increase in apoptosis, q_p_ increased across all LgEV treatment conditions, reaching 35 pg/cell/day, a 3.5-fold increase compared to the control.

### 3.3 Role of glutamine in rescuing the impact of LgEVs

Glutamine supplementation (**Gln^+^**) effectively mitigated the growth-inhibitory and pro-apoptotic effects of LgEVs while preserving their positive effect on specific productivity (Figures 2 and 3). In Gln^+^ cultures treated with fed-batch-derived LgEVs (2–4 × 10⁷ LgEVs/mL), the growth inhibition observed in Gln^-^ conditions was largely reversed. Under these conditions, we detected only a marginal increase in early apoptosis at the highest dose (6 × 10⁷ LgEVs/mL), from 1.5% to 2% (Figure 3B). We observed a similar protective trend with LgEVs from Day 11 perfusion cultures. In Gln^+^ conditions, addition of 2 × 10⁷ and 4 × 10⁷ LgEVs/mL reduced VCD by 17% and 37%, respectively, by day 4 (Figure 3E). In contrast, VCD reductions in Gln^-^ conditions reached 64.4% and 90% for the same dose, respectively. Apoptosis assays further confirmed that Gln attenuated the apoptotic effect of perfusion LgEVs. At the higher dose of 4 × 10⁷ LgEVs/mL, the late-apoptotic population in Gln^+^ cultures was only 7.2%, compared to 25.4% in Gln^-^ cultures—a 3.5-fold reduction in apoptotic burden. Notably, Gln^+^ cultures still exhibited high mAb productivity with EV treatment. These results indicate that glutamine availability decouples the pro-productivity signals carried by LgEVs from their detrimental impact on cell proliferation and survival.

**Figure 3.**
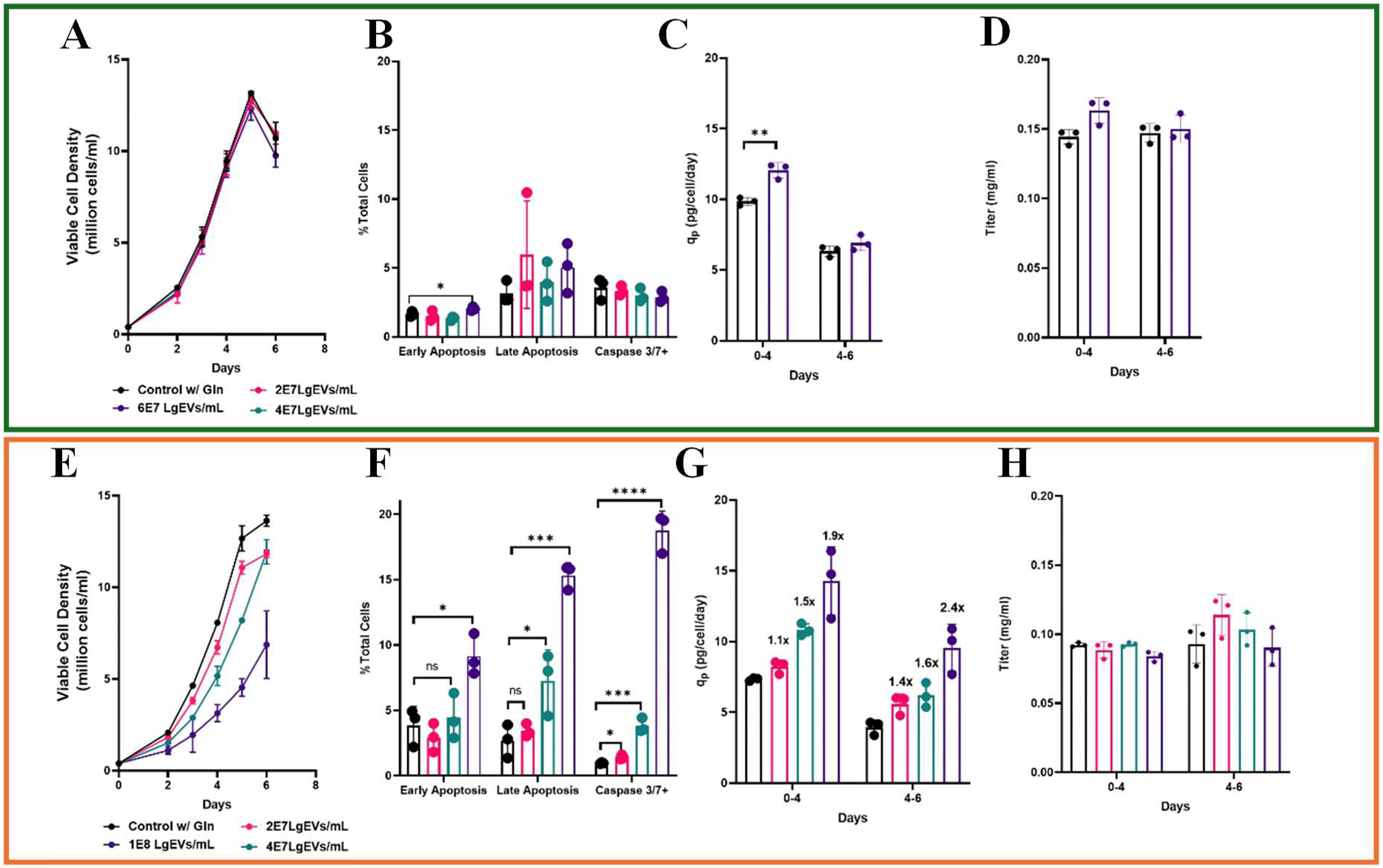
Impact of different dosages of LgEVs from D8 fed-batch cultures (panels A-D) and D11 perfusion culture (panels E-H) on cell growth, productivity, and apoptosis of fresh CHO cultures with glutamine addition (Gln^+^). (A-D): Gln^+^ cultures after adding D8 fed-batch LgEVs (2×10^7^, 4×10^7^, 6×10^7^ LgEVs/mL) or no EVs (control). (E–H) Gln^+^ cultures after adding D11 perfusion LgEVs (2×10^7^, 4×10^7^, 1×10^8^ LgEVs/mL) or no EVs (control). (A, F) Viable cell density (million cells/ mL) over 6 days after adding EVs on day 0. (B, F) Percentage of early and late apoptotic cells at day 4 of cultures as assessed by AnnexinV/7AAD, and by Caspase 3/7 flow cytometry assays. (C, G) Cumulative Antibody titer (mg/ml) from day 0 to day 4 and from day 4 to day 6. (D, H) Cell-specific antibody productivities (q_p_): day 0 to day 4 and day 4 to day 6 (pg/cell/day). Error bars represent standard deviation of biological replicates (n=3).

### 3.4 LgEVs alter CHO cell amino acid and sugar metabolism but differently in Gln^+^ versus Gln^-^ conditions

To understand the metabolic basis of the observed phenotypic shifts in perfusion LgEV-treated cells, we measured the consumption and production rates of amino acids, glucose, and lactate between day 2 and day 4 post-treatment (Figure 4). While LgEVs addition increased baseline glucose consumption in both environments, the metabolism of a unique set of amino acids (notably alanine, glycine, serine, and asparagine) was heavily modulated by Gln availability and LgEVs dosage.

**Figure 4.**
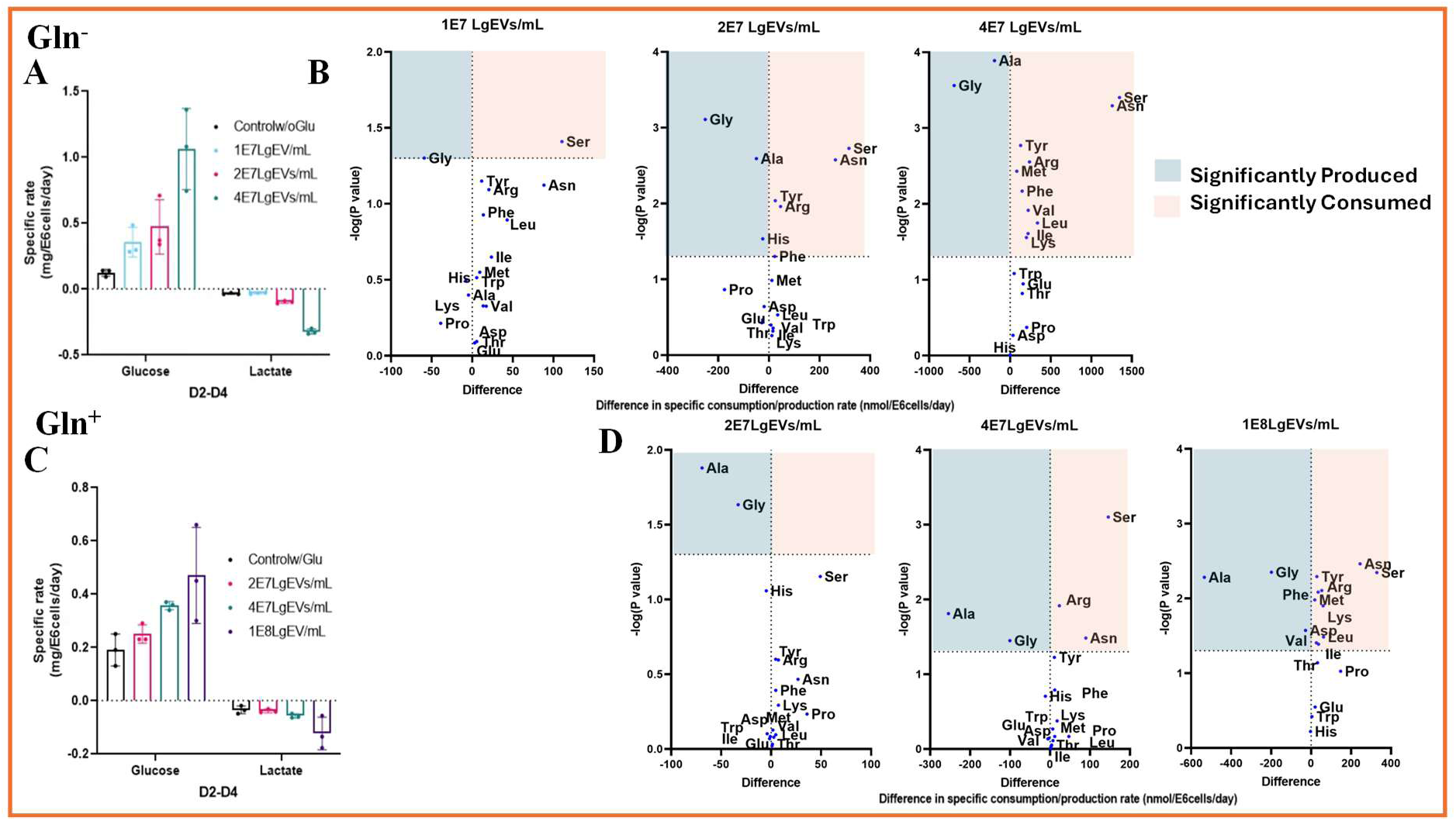
Glucose and amino acid utilization and lactate production from Day 2 to Day 4 in CHO cells treated with D11 Perfusion LgEVs with (Gln^+^) and without glutamine (Gln^-^) addition compared to controls. **(A–B) Gln^-^** cultures treated with different concentrations of LgEVs (1×10^7^, 2×10^7^, 4×10^7^) or no EVs (control). (C-D) **Gln^+^** cultures treated with LgEVs (2×10^7^, 4×10^7^, 1×10^8^ LgEVs/mL) or no EVs (control). **(A,C):** Significant increase in specific glucose consumption and lactate production rates (mg/10⁶ cells/day) in response to increasing doses of LgEVs. **(B)** Volcano plots show differential amino acid consumption and production rates (nmol/10⁶ cells/day) in cells treated with 1×10^7^, 2×10^7^, 4×10^7^ LgEVs/mL compared to **Gln^-^** control. **(D)** Volcano plots show differential amino acid consumption and production rates (nmol/10⁶ cells/day) in cells treated with 2×10^7^, 4×10^7^, 1×10^8^ LgEVs/mL compared to **Gln^+^** control. Amino acids above the horizontal dotted line are significantly altered (*p* < 0.05). Amino acids to the right of the vertical line are consumed at higher rates, while those to the left are produced more compared to control.

Under Gln^-^ conditions, the lowest dose of EVs (1×10^7^ LgEVs/mL) tripled the specific glucose consumption rate relative to the control without significantly increasing lactate secretion. Volcano plot analysis revealed that at the low dose, changes in amino acid metabolism were largely confined to accelerated consumption of serine and asparagine alongside a mild increase in glycine secretion (Figure 4B, left panel). However, as the LgEV dose increased to 2 × 10⁷ and 4 × 10⁷ LgEVs/mL, a pronounced, dose-dependent metabolic overflow emerged. The mid LgEVs dose maintained high glucose uptake while triggering a marked increase in lactate secretion (from 0.03 mg/10^6^ cells/day to 0.1 mg/10^6^ cells/day). On the other hand, the highest LgEV dose triggered a massive surge in lactate secretion, pushing the specific rates of glucose consumption and lactate secretion up to 10-fold higher (Figure 4A). The shift indicates that under the stress of high LgEVs concentrations, Gln^-^ cells could not fully utilize pyruvate (from glucose glycolysis) into the TCA cycle, instead apparently diverting it to form lactate. Concurrently, more amino acid uptake rates reached statistical significance, starting with serine and asparagine at the middle dose (Figure 4B, middle and right panels). As EVs dosage increased, more branched amino acids (valine, leucine, isoleucine), and the aromatic amino acid tyrosine consumption rates increased dramatically.

Addition of LgEVs in the Gln^+^ environment yielded a somewhat similar but distinct metabolic pattern compared to the Gln^-^ condition. Here, glucose consumption increased more modestly with EV addition, and lactate secretion remained stable and low across all doses (Figure 4C). These results show that cells do not have to rely heavily on glycolysis when Gln is present. The most significant trend at 2×107 LgEVs/mL was an increase in alanine and glycine secretion relative to the control.

Direct comparison between these two cultures (Gln^+^ vs Gln^-^) at the same LgEV dosage highlights the clear impact of Gln availability on EV processing. A notable difference appeared in alanine and glycine secretion patterns under both 2×10^7^ LgEVs/mL and 4×10^7^ LgEVs/mL doses (Figure 4B vs. 4D; & Figure 5). In Gln^-^ cultures, glycine was the dominant secreted amino acid, scaling with EV dose and shifting far to the left on the volcano plot (reaching nearly 600 nmol/10^6^ cells/day in Figure 4B, right), with alanine production barely statistically significant (Figure 5). Conversely, in Gln^+^ cultures, this relationship reversed, with alanine becoming the dominant secreted amino acid and glycine secretion strongly suppressed (Figure 4D, Figure 5). At the same time, crucial amino-acid carbon sources such as asparagine, serine, and arginine were significantly increased in the absence of Gln.

**Figure 5.**
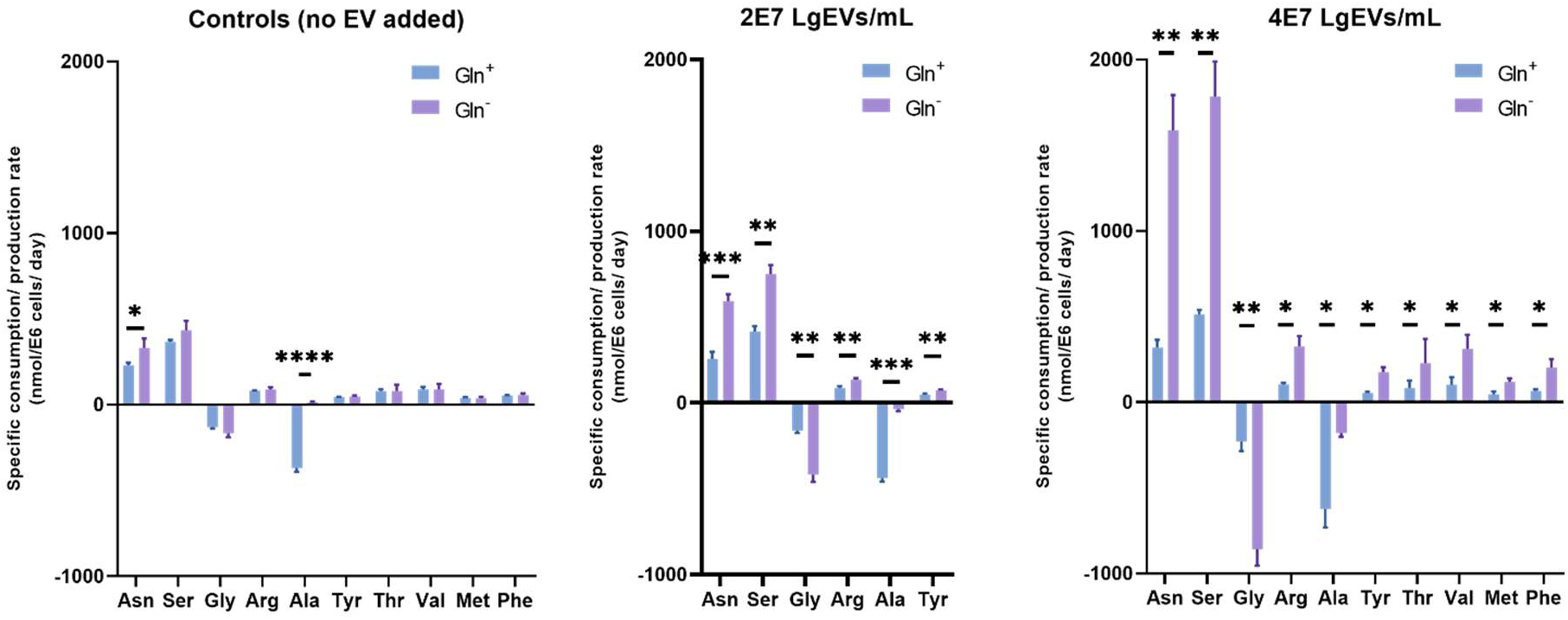
Specific amino acid consumption and production rates from day 2 to day 4 at two dosages of LgEVs (from D11 perfusion culture) in Gln^+^ and Gln^-^ cultures (n=3). (A): Controls (no EVs added). (B): 2 × 10⁷ LgEVs/mL added. (C) 4 × 10⁷ LgEVs/mL. in Panel B and C, we plotted only amino acid that showed statistically significant differences. Panel A contains all amino acids that are listed in panels B and/or C.

### 3.5 Metabolic flux analysis (MFA) provides insights into the quantitative intracellular flux changes due to LgEV addition

To quantify intracellular changes in carbon distribution, we estimated metabolic fluxes from day 2 to day 4 post-treatment (experiments corresponding to Figures 4 and 5) using a validated, compact stoichiometric model of CHO cell metabolism (Venkatarama Reddy et al., 2026). Analyzing the distinct shifts in flux distributions between control and LgEV-treated cultures reveals a highly coordinated metabolic reconfiguration. Overall, late-stage LgEVs induce a metabolically stressed phenotype characterized by a severe bottleneck at the pyruvate node, an increased reliance on amino acid-supported anaplerosis, a glutamine-dependent switch between glycine and alanine overflow metabolism, and a strategic redistribution of resources away from proliferation toward maintenance and antibody synthesis.

Arrow colors represent flux changes; blue: flux increase; green: flux decrease; red: flux reversal, while arrow thickness scales with the magnitude of change (thin: negligible change; number of % change are shown on select fluxes).

Notes: 1) Certain pathways, such as the interconversion of alanine, pyruvate, glutamate, and aKG can occur concurrently across both the cytoplasm and mitochondrial compartments; 2) Not all the fluxes of Supplemental Excel Table 1 are shown in this graph.

**Table 1.**
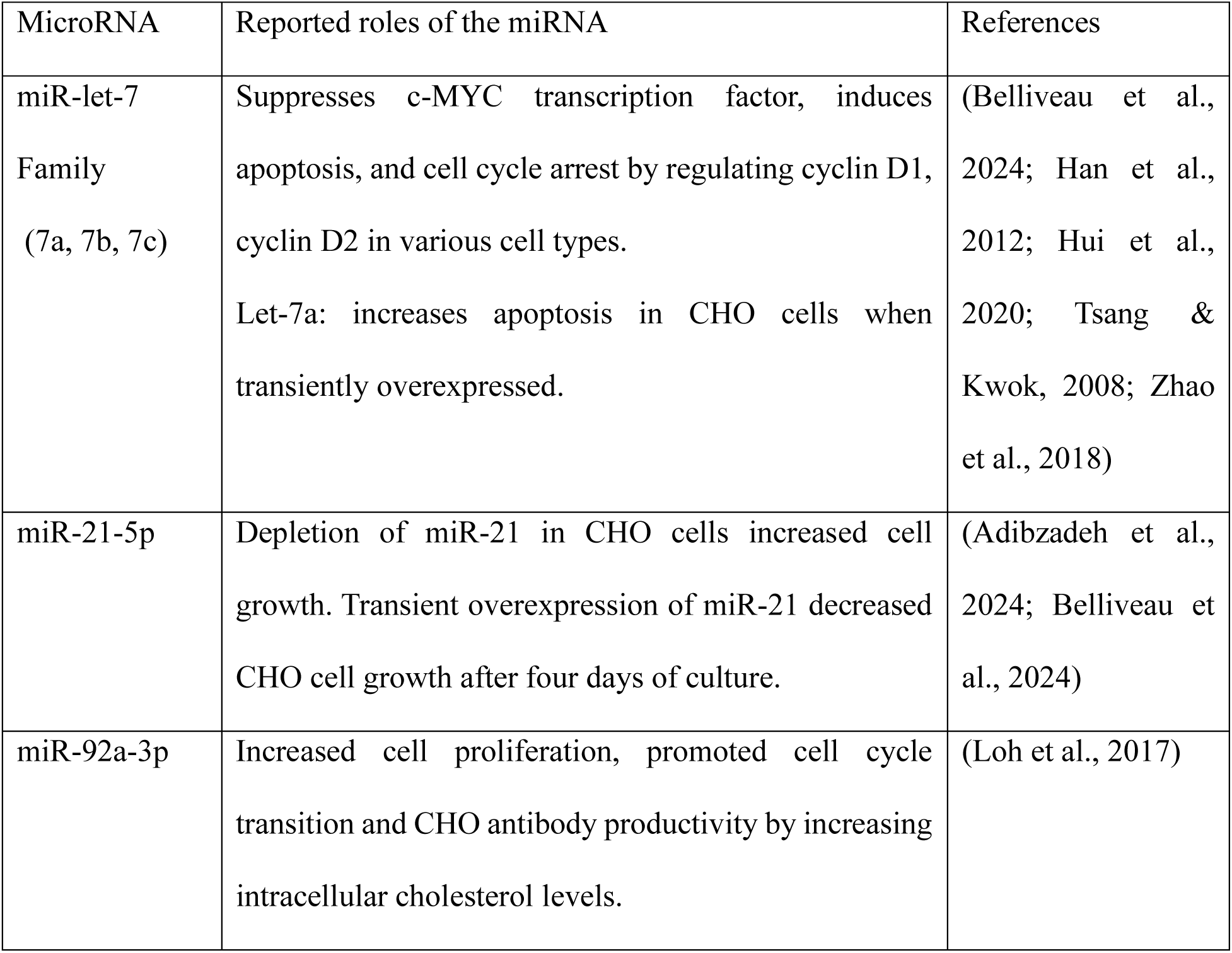
List of miRs and their reported roles

#### 3.5.1 A metabolic bottleneck at the pyruvate node drives lactate overflow

A notable consequence of LgEV treatment, particularly in Gln^-^ conditions, was the emergence of a severe metabolic bottleneck at the pyruvate node. Under these conditions, the glucose-6-phosphate (GAP) flux from glucose surged by 125% relative to the control (Figure 6). However, this glycolytic carbon flux could not be efficiently funneled into the TCA cycle. In contrast, the flux of pyruvate into the TCA cycle increased by a marginal 8.9%, and the flux from pyruvate to lactate expanded by 274% (Figure 6). This flux partitioning indicates that the cells reached a functional capacity limit at the pyruvate node, driving a massive metabolic overflow toward lactate secretion.

**Figure 6.**
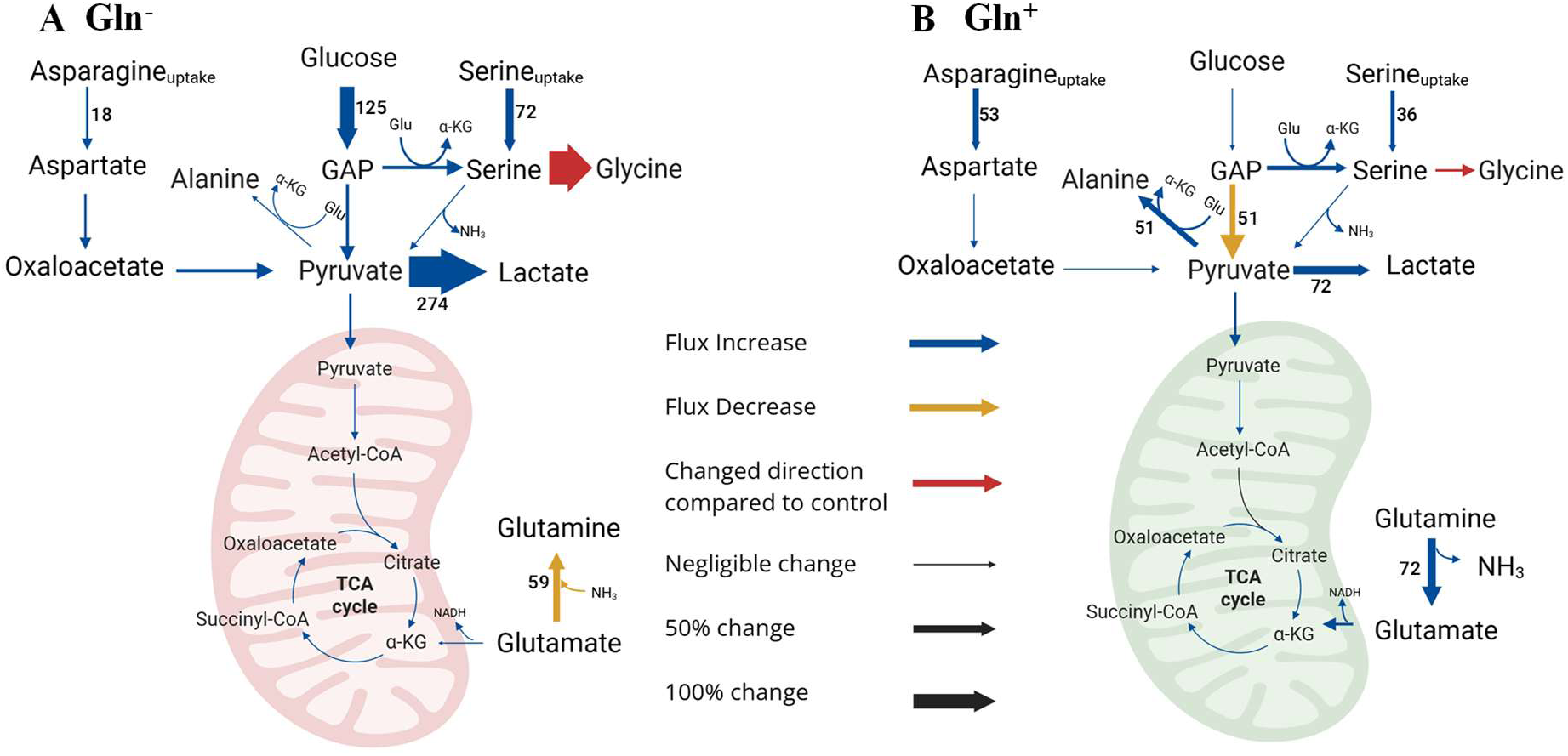
Metabolic Flux Analysis (MFA) of CHO cells under LgEV treatment conditions. Percent MFA flux differences in CHO cells between LgEV-treated and controls groups. (A): **Gln^-^**cultures with 2×10^7^ LgEVs/mL addition relative to control (no EVs added). (B): **Gln^+^** cultures with 4×10^7^ LgEVs/mL addition relative to control (no EVs added).

#### 3.5.2 Recruitment of alternative anaplerotic substrates

In Gln^-^ cultures, to compensate for the lack of glutamine-driven anaplerosis, EV-treated cultures increasingly recruited alternative anaplerotic substrates, namely branched-chain amino acids (BCAAs), with intracellular fluxes for valine and leucine catabolism increasing by 20% and 33%, respectively (Supplemental Excel Table 1). This activation indicates an attempt to alleviate energetic stress by oxidizing BCAAs into key TCA cycle intermediates (Leucine → acetyl-CoA; Valine → succinyl-CoA). Additionally, asparagine-to-aspartate conversion rose by 18% in Gln^-^conditions and by 53% in Gln^+^ conditions, routing alternative carbon pools into oxaloacetate and pyruvate to support the network (Figure 6, Supplemental Excel Table 1).

#### 3.5.3 The glutamine-dependent glycine and alanine secretion switch

One of the most striking metabolic signatures associated with EV exposure was the distinct switch from high glycine secretion in Gln^-^ cultures to high alanine secretion in Gln^+^ cultures (Figure 4,5, and 6). In Gln^+^ cultures, LgEV treatment triggered a 72% increase in glutamine deamination to glutamate, which subsequently fueled the α-ketoglutarate (αKG) pool (Figure 6 and Supplemental Excel Table 1). This EV-induced glutamine hyperconsumption aligns with reconfigurations observed in cancer biology, in which stressed cells prioritize glutamine-driven anaplerosis (Zhao et al., 2016). The resulting accumulation of intracellular pyruvate, combined with increased αKG driven by LgEV treatment, led to a 51% increase in the alanine transamination flux in Gln^+^ cultures (Figure 6). This marks a classic alanine transamination overflow, a hallmark of glutamine-supported mammalian metabolism (Kirsch et al., 2022; Chevallier et al., 2020). In Gln^-^ cultures, lacking glutamine as a nitrogen donor, the cells compensated by upregulating serine uptake by 72% and channeling the carbon toward glycine production via serine-glycine one-carbon metabolism with up to 3.5-fold increase in magnitude compared to control (Figure 6 and Supplemental Excel Table 1). In both Gln^+^ and Gln^-^ cultures, the formation of glycine from serine constitutes a flux reversal compared to control (no EVs added).

#### 3.5.4 Elevated energy demands and glutamine-mediated protection

Despite differences in glutamine availability, the addition of LgEV imposed a consistent energetic signature across both experimental sets. We observed a marked increase in the oxidation of FADH_2_ to generate ATP, showing a 33% increase in Gln^-^ cultures and a 29% increase in Gln^+^ cultures treated with LgEV compared to their respective controls (Supplemental Excel Table 1). This elevated requirement for FADH_2_ generation indicates an increased demand for ATP under EV-associated stresses, such as vesicle processing or hyperactivated protein synthesis.

Indeed, we hypothesized that glutamine could rescue LgEVs-treated cells from apoptosis by providing abundant ATP. To test this hypothesis, we performed MFA on the first 3 days of baseline batch cultures ***without LgEVs addition*** with and without glutamine supplementation (Figure S3). MFA profiles showed that initial glutamine supplementation significantly bolsters the baseline pool of electron carriers, resulting in 68% higher FADH2 and 46.7% higher total NADH formation fluxes during the first two days of culture (Figure S3). This offers a clear explanation for the glutamine rescue effect: the pre-existing high-energy supply from glutamine supplementation provides cells with an ATP cushion. This additional energy capacity allows the cells to tolerate late-stage EV stressors, effectively preventing the severe ATP deficit (ATP demand > ATP generation) that otherwise drives cells in Gln^-^ cultures into apoptosis.

#### 3.5.5 Metabolic reallocation: driving the growth–productivity tradeoff

Collectively, the structural burden of the pyruvate bottleneck, energetic stress, and amino acid catabolism suppress biomass accumulation, thereby slowing cell growth. By curtailing cell division, the cell drastically reduces its metabolic demand for biomass synthesis. This reduction frees up a significant pool of high-value ATP and cellular machinery. Consequently, the cell undergoes a strategic metabolic reallocation, directing these newly available resources away from proliferation and toward cellular maintenance and high-intensity recombinant protein secretion. This systemic shift ultimately drives the observed dose-dependent boost in cell-specific mAb productivity.

### 3.6 Removal of LgEVs enhances CHO-cell growth thus confirming the negative impact of LgEVs on cell performance

To further investigate the effects of different EV subpopulations accumulating in perfusion cultures, we conducted “satellite” batch cultures using a sequential media-fractionation strategy. Our goal was to isolate the phenotypic impact of each component of the used perfusion-culture medium by selectively removing apoptotic bodies, LgEVs, sEVs, large proteins and particles (100 kDa), and small proteins (5 kDa) from day 7 perfusion spent media. Then we resuspended the conditioned cells (from day 7 perfusion) in the various media fractions. We hypothesized that removal of detrimental LgEVs would improve cell growth. As shown in Figure 7, on day 7 of the perfusion process, cells in media fractions without apoptotic bodies (ApBFree) and LgEVs free (LgEVFree) grew better than the control from day 8 onward. In particular, the LgEVFree cultures can sustain viable-cell growth longer, achieving a 4-fold higher VCD at day 11 compared to the control. Surprisingly, although cell growth increased, the antibody titer of LgEV-free cultures remained unchanged compared to the control.

**Figure 7.**
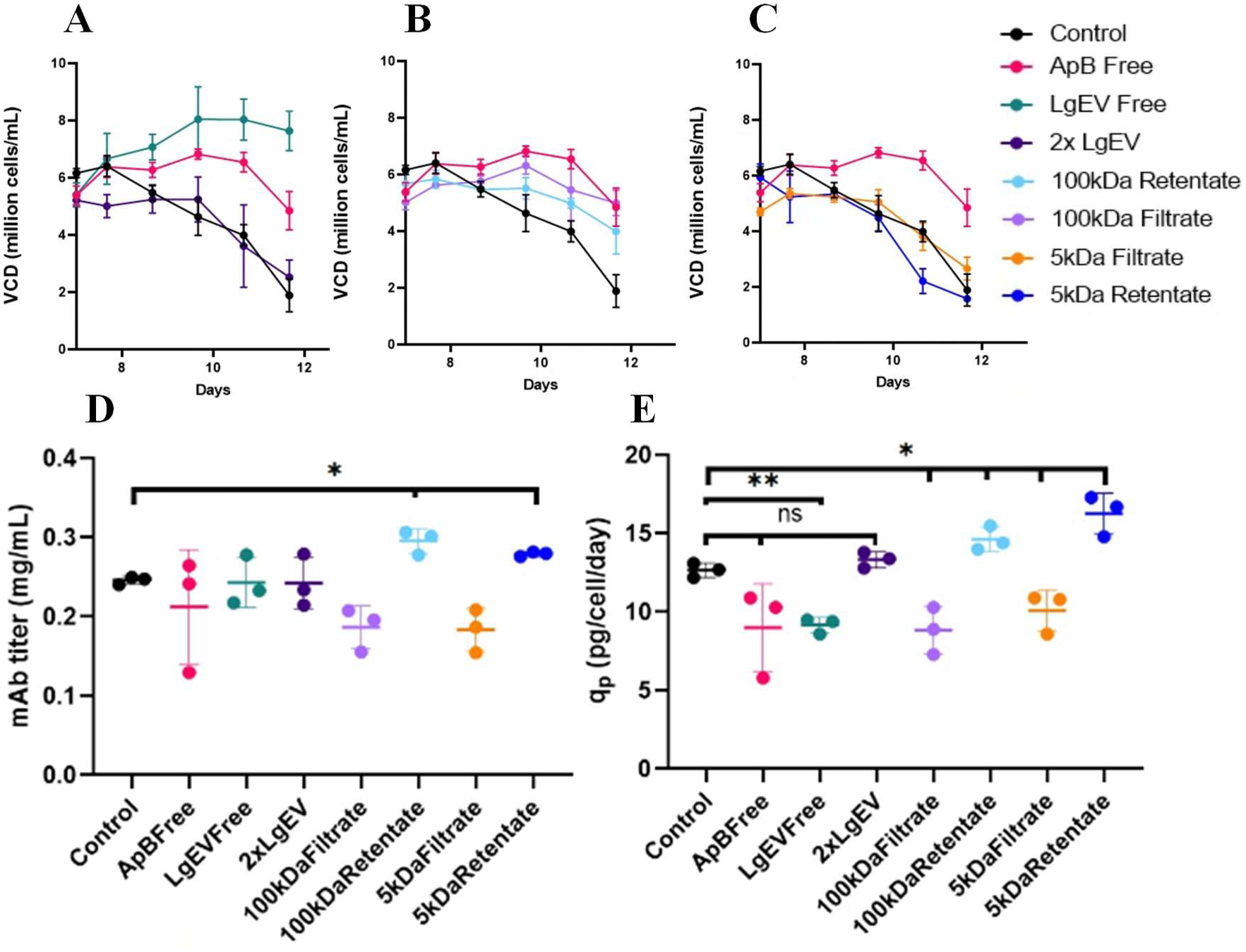
Cell growth and productivity of “satellite” cultures in various media fractions on day 7 of perfusion cultures. (A-C) The viable cell density (million cells/ mL) over time for cultures under eight experimental conditions: Control, Apoptotic bodies free (ApB Free), LgEV Free, 2× LgEV, 100 kDa Retentate, 100 kDa Filtrate, 5 kDa Filtrate, and 5 kDa Retentate. (D) Cumulative Antibody titer: (mg/mL) from day 0 to day 4 and from day 7 to day 11 (E) cell-specific antibody productivity (q_p_) across the same conditions from day 7 to day 11 (pg/cell/day).

In bioprocess applications, EVs have been an overlooked component in assessing different perfusion cell retention device (CRD) technologies. Table S1 summarizes common CRDs and their expected capacity to remove EVs. While size-based separation methods (ATF, TFF) face limitations due to size overlap between EVs and membrane pore sizes, density-based separation methods (such as acoustic wave separators or centrifuges) could selectively clear these populations. Because current filtration systems often face fouling difficulties during long, high-density perfusion runs, optimizing CRD selection for targeted EV removal represents a promising strategy to mitigate the performance drops observed in late-stage cultures (Zhang et al., 2024).

In contrast to the perfusion results, removing EVs from fed-batch cultures yielded highly variable outcomes and were not statistically significantly different (data not shown). This variability is likely due to fed-batch cells being more metabolically exhausted and already transitioning toward G1-phase arrest by day 7 (Figure S4). Consequently, once the cell population is committed to the G1 non-dividing state, removing EVs is no longer sufficient to restore growth, making the window for effective EV removal much narrower in fed-batch cultures compared to perfusion.

### 3.7 Can the enrichment of a subset of miRNAs explain the impact of EV on cell growth and metabolism?

To investigate whether the biological activity of late-stage LgEVs may be associated with their miRNA cargo, we used RT-qPCR assays to quantify five selected miRNAs in parent cells and isolated LgEVs harvested from D5 and D8 fed-batch cultures (Figure 8). These miRNAs were selected based on their abundance in CHO cells and EVs, and their previously reported roles in regulating proliferation, stress adaptation, apoptosis, and productivity (Belliveau & Papoutsakis, 2023; Belliveau et al., 2024).

**Figure 8.**
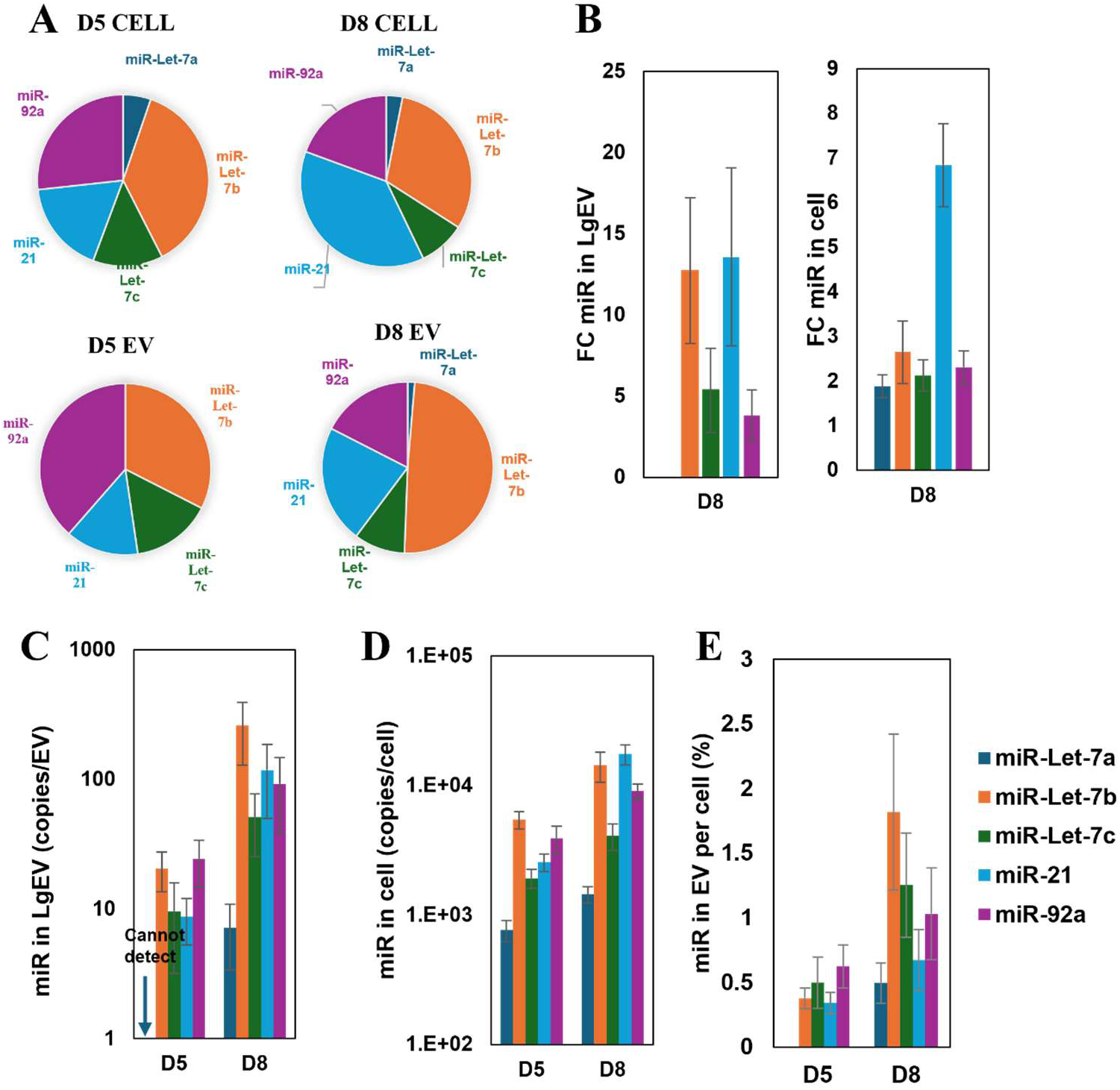
Profiling selected miRNAs in CHO cells and their LgEVs at D5 and D8 of fed-batch cultures. (A) Pie charts depict the relative composition of selected miRNAs (miR-Let-7a, miR-Let-7b, miR-Let-7c, miR-21, and miR-92a) in EVs and cells at D5 and D8. (B) Fold change plots represent the relative enrichment of each miRNA in EVs and cells at D8 normalized to D5. Error bars represent the propagated standard deviation. (C-D) Bar plots show the absolute quantification of miRNA copies per LqEV (C) and per cell (D). Error bars represent the standard deviation of three biological replicates. (E) Ratio of each miRNA copies found in EV versus cell (%). Error bars represent the propagated standard deviation.

RNAseq analysis of the CHO microRNome previously showed that among more than 600 detectable miRNAs, five miRNAs account for approximately one-half of the total cellular and EV-associated miRNA content. Importantly, as we have previously reported, while unstressed CHO cultures are characterized by high abundances of miR-92a and miR-23a, ammonia and osmotic stress induce a marked enrichment of let-7 miRNA family members, particularly let-7c, let-7b, and let-7a, in both cells and EVs (Belliveau & Papoutsakis, 2023). In the same study, the let-7 family was proposed to constitute a conserved stress-responsive regulatory module in CHO cells. Subsequently, using stoichiometric RT-qPCR analyses, we have demonstrated selective loading of these abundant miRNAs into CHO EVs and showed that overexpression of let-7a decreases CHO-cell viability, whereas overexpression of miR-92a promotes cell survival; miR-21 overexpression exerts a modest detrimental effect on growth and viability during late exponential culture (Belliveau et al., 2024). Of note, although the let-7 miRNA family members (let-7a, let-7b, and let-7c) are reported separately, they all target the same set of genes through the same seed sequence (GAGGUAG).

Consistent with these previous observations, here, we found substantial enrichment of let-7b, let-7c, and miR-21 in D8 LgEVs compared with D5 vesicles, with copy numbers increasing approximately 5–15-fold on a per-vesicle basis. These D8 vesicles also elicited the strongest phenotypic responses in recipient cultures, including inhibition of proliferation, increased apoptosis under glutamine limitation, and enhanced cell-specific mAb productivity. The observed phenotypes are remarkably consistent with the known biological activities of the let-7 family, which regulates proliferation, differentiation and stress adaptation through targets that include HMGA2, RAS, MYC, CDK6 and components of the IGF signaling pathway (Roush & Slack, 2008; Zhu et al., 2011), as well as with the reported effects of miR-21 and miR-92a on CHO-cell growth and survival (Belliveau et al., 2024). We have previously demonstrated that the CHO EVs mediate massive exchange of RNA and proteins between cells in CHO cultures (Belliveau & Papoutsakis, 2022), and separately, in a different cell-EV system, that LgEVs (formerly referred to as microparticles, MPs) effectively deliver RNA cargo to target cells and that the cargo produces the expected regulatory phenotype (Kao & Papoutsakis, 2018). Taken together, these data support the effective delivery of LgEV functional cargo (including miRNAs) to CHO cells.

## IV Conclusions

This study sought to explore an overlooked component in spent media, namely extracellular vesicles. We profiled EVs and their potential activities in CHO batch, fed-batch, and perfusion cultures. We found that EVs could act as a dual mediator of CHO-cell growth and productivity. Specifically, LgEVs from day 8 fed-batch and day 11 perfusion cultures reduced cell growth but increased cell-specific productivity. A subset of miRs was profiled in fed-batch EVs to show their enrichment profile, which was distinct from that of their parent cells, thus demonstrating active miR sorting by the CHO cells. Our data show that EVs in CHO cell culture could alter amino acid metabolism. We identified key differences when EVs were added to culture with and without Gln. Lastly, we attempted to mitigate their effects via media fractionation and produced a list of CRDs and their potential for EV removal.

These results underscore the need for further investigation into EVs generated under industrially relevant conditions, such as after biphasic operations, under ammonia-stress conditions, or in other CHO cell lines. Future work could aim to gain deeper mechanistic insight through proteomics and RNA-Seq analyses of the subset of EVs that affect cell growth and productivity, to identify the specific proteins and miRNAs responsible for the observed effects. This approach could reveal the specific miRNAs released under different types of stress, serving as signatures for monitoring cell culture health, especially since various stresses can manifest as common symptoms such as reduced growth and elevated lactate secretion. It can also suggest potential targets for cell-line engineering of EV-associated molecules to optimize bioprocess outcomes.

One of the strongest precedents for EV-mediated metabolic remodeling was reported by Zhao et al. (2016), who demonstrated that sEVs (exosomes) derived from cancer-associated fibroblasts (CAFs) suppress oxidative phosphorylation and promote glycolysis in prostate and pancreatic cancer cells. Importantly, CAF-derived exosomes induced glutamine-dependent reductive carboxylation and supplied recipient cells with metabolites capable of supporting survival and biosynthesis under nutrient-limiting conditions (Zhao et al., 2016). Subsequently, Achreja et al. (Achreja et al., 2017) developed Exo-MFA, a ^13^C-metabolic flux analysis framework specifically designed to quantify intracellular flux redistribution resulting from exosome-mediated metabolite transfer. The findings presented here exhibit notable conceptual similarities to these cancer studies. In both systems, EV exposure is associated with increased glycolytic activity and significant remodeling of glutamine and TCA-cycle metabolism. However, unlike CAF-derived exosomes, which predominantly support tumor cell growth and survival, late-stage CHO-cell LgEVs appear to transmit a stress-associated metabolic program characterized by growth inhibition, increased apoptosis under glutamine limitation, enhanced glycolytic overflow, altered amino acid utilization, and increased cell-specific productivity. Glutamine supplementation largely alleviated the detrimental effects of LgEVs on growth and viability while preserving enhanced productivity, suggesting that glutamine availability modulates how recipient CHO cells process EV-induced metabolic stress. Given that CHO cells exhibit many hallmarks of transformed mammalian cells, including aerobic glycolysis and glutamine addiction, the parallels between EV-mediated metabolic remodeling in cancer cells and CHO cultures are particularly intriguing and warrant further investigation.

## V. Credit authorship contribution statement

**H.B.N.:** Conceptualization, Data curation, Formal analysis, Investigation, Methodology, Visualization, Writing – original draft, Writing – review & editing. **N.G.M:** Data collection, Investigation, Software, Writing – review & editing. **A.P**: Investigation. **K.H.L.:** Resources, Supervision, Writing – review & editing. **E.T.P.:** Conceptualization, Data curation, Funding acquisition, Project administration, Resources, Supervision, Writing – review & editing.

## Acknowledgments

This work was funded by the U.S. Food and Drug Administration (FDA), under Award Number U01FD007695, and the Advanced Mammalian Biomanufacturing Innovation Center (AMBIC), under Award Number 2005913780 and parent AMBIC award number NSF EEC-2100502.

## VI. Nomenclature

- 7AAD: 7-aminoactinomycin D
- AKG: alpha-ketoglutarate
- ApB: Apoptotic body
- ATF: Alternating tangential flow
- ATP: Adenosine triphosphate
- BCAA: Branched-chain amino acid
- CAF: Cancer-associated fibroblast
- CHO: Chinese Hamster Ovary
- CRD: Cell retention device
- DO: Dissolved oxygen
- EV: Extracellular vesicle
- FADH_2_: Flavin adenine dinucleotide
- GAP: Glucose-6-phosphate
- Gln: Glutamine
- HPLC: High-performance liquid chromatography
- IgG: Immunoglobulin G
- ISEV: International Society for Extracellular Vesicles
- IVCD: Integral viable cell density
- LgEV: Large extracellular vesicle
- lncRNA: Long non-coding RNA
- mAb: Monoclonal antibody
- MFA: Metabolic flux analysis
- miRNA / miR: MicroRNA
- MWCO: Molecular weight cut-off
- NADH: Nicotinamide adenine dinucleotide (reduced)
- NTA: Nanoparticle tracking analysis
- PBS: Phosphate-buffered saline
- PES: Polyethersulfone
- piRNA: Piwi-interacting RNA
- q_M_: Specific metabolite consumption/production rate
- q_p_: Cell-specific antibody productivity
- RT-qPCR: Reverse transcription quantitative polymerase chain reaction
- sEV: Small extracellular vesicle
- TCA: Tricarboxylic acid
- TFF: Tangential flow filtration
- VCD: Viable cell density
- VSSC: Violet side scatter mode
- vvd: Vessel volumes per day

## Data Availability Statement

Data will be made available upon request.

## Declaration of Generative AI and AI-assisted technologies in the writing process

The authors used ChatGPT (OpenAI) and Gemini (Google) as language-assistance tools to improve the clarity and readability of portions of the manuscript. All scientific content, analyses, and conclusions were developed, verified, and approved by the authors.

