## Supplemental Materials for "Late-Stage Large Extracellular Vesicles Reprogram CHO Cell Metabolism in a Glutamine-Dependent Mode and Promote Antibody-Productivity to Cell-Growth Tradeoff"

### Supplementary Figures

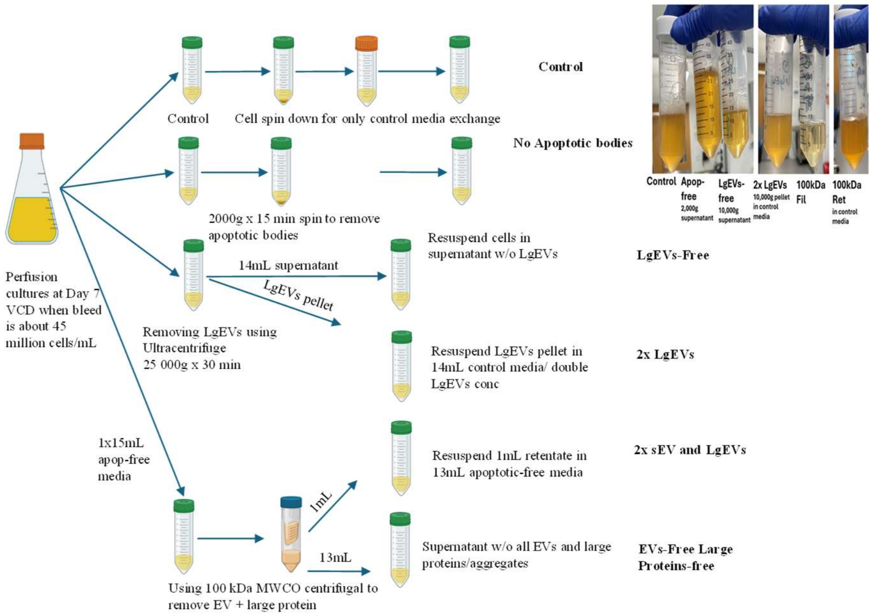


Figure S1. Experimental design of fed-batch and perfusion satellite cultures.

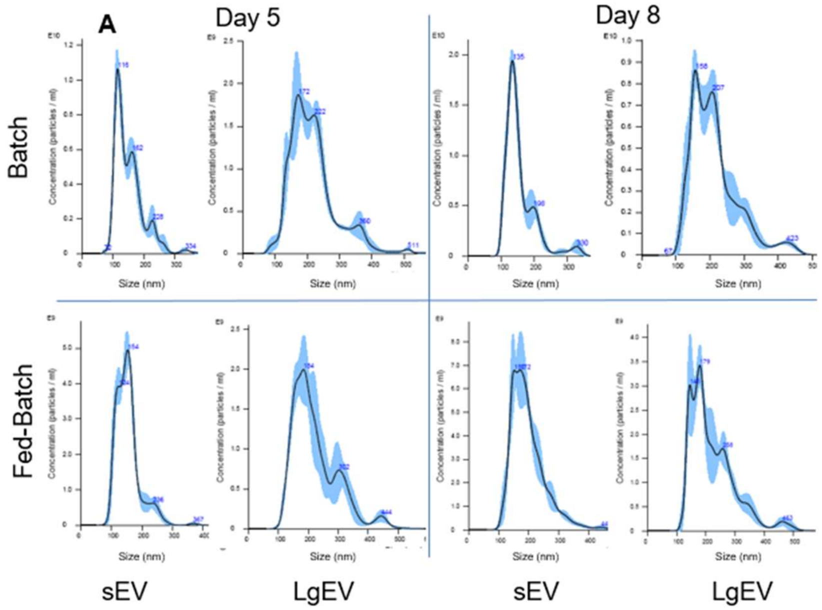


Figure S2. Nanoparticle tracking analysis of LgEVs and sEVs from D5 and D8 of fed-batch cultures.

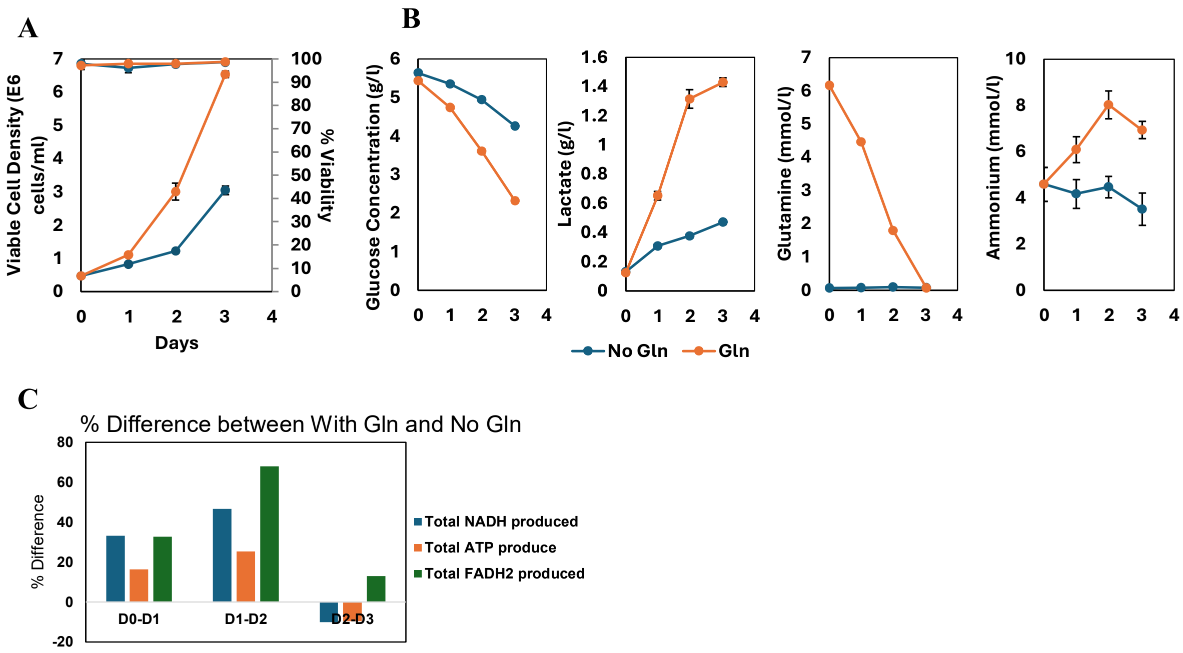


**Figure S3. Short term batch cultures of CHO cells.** (A) Growth profiles showing viable cell density (VCD) and viability (%) for $Gln^+$ (6 mM Gln supplementation) and $Gln^-$ (no Gln) cultures. (B) Metabolite profiles for glucose, lactate, glutamine, and ammonium highlighting the rapid depletion of glutamine by Day 3 and the shift toward higher lactate secretion in $Gln^+$ cultures. N = 3 (C) Differential energy carrier production (NADH, ATP, $FADH_2$) calculated from MFA analysis and plotted as % difference between $Gln^+$ and $Gln^-$ cultures.

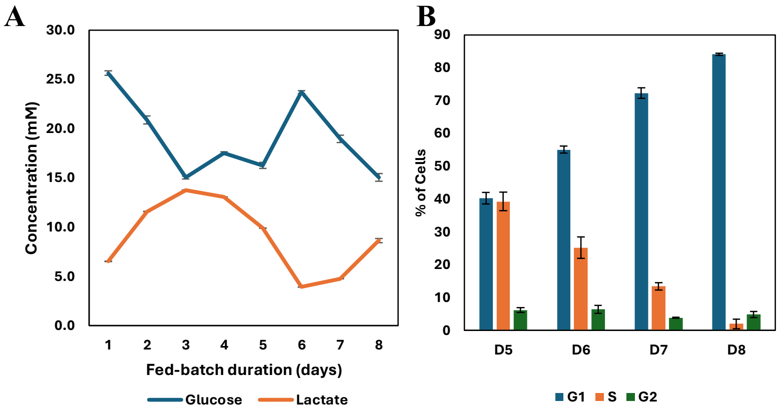


**Figure S4. Standard fed-batch CHO cultures with initial 6mM Gln supplementation where EVs were collected from.** (A) Residual glucose and lactate concentration profiles on each day prior to feeding. (B) Cell cycle distribution of cells in D5, D6, D7 and D8 via the EdU assay (n=3).

Table S1. Current cell retention device options and their potential for removing EVs

| | Cell Retention Device | Description | Can it hypothetically remove EVs? | References |
|---|---|---|---|---|
| Retention by particle size: Filtration-based | Hollow Fiber: tangential flow filtration (TFF), alternating tangential flow (ATF) | Applying a high liquid flow rate at low shear at the membrane surface to harvest spent medium | Depending on filter pore size, most reported studies used 0.2-0.65 μm but the effective pore size decreases over time due to membrane fouling | (Clincke et al., 2013; Voisard et al., 2003; Zhang et al., 2024) |
| | Spin-filter | Internal or external spin filters with a wire screen rotate, retaining cells due to the steric effect of the rotating screen | Potentially yes, depending on screen materials and filter pore size. Pore sizes are usually larger than the average cell size (10, 20, 25 μm). Polymer screens are less prone to fouling and clogging | (Esclade et al., 1991; Himmelfarb et al., 1969) |
| Retention by particle density | Centrifuge | Using centrifugal force | Yes, because EVs do not settle at 300xg | (Welsh et al., 2024) |
| | Acoustic waves enhanced settler | Applied acoustic resonance field to separate cells, can have a saturation limit | Highly yes. A study demonstrated superior performance using acoustic settler over ATF for influenza virion production, successful continuous harvest of virion also | (Granicher et al., 2020; Trampler et al., 1994) |

| | | | | |
|---|---|---|---|---|
| | | | demonstrate ability to remove EVs | |
| | Gravity Settlers: Inclined settler | Via the Boycott effect, cells settle more efficiently on an inclined surface and slide down to be retained in the reactor | Yes, because EV density is much lower than cell density.<br>Research showed it was more efficient at retaining viable than dead cells (Batt et al., 1990), hence EV could also be removed with with dead cells | (Batt et al., 1990; Voisard et al., 2003) |
| | Hydrocyclone | Cyclonic separators use differences in density of feed and centrifugal movement by injecting feed tangentially to the wall | Unsure. However, loss of viability from 5 to 12% makes it not ideal for mammalian CRD | (Jockwer et al., 2001) |
