## Supplemental Excel Table 1 for "Late-Stage Large Extracellular Vesicles Reprogram CHO Cell Metabolism in a Glutamine-Dependent Mode and Promote Antibody-Productivity to Cell-Growth Tradeoff"

Average experimental and Metabolic Flux Analysis (MFA) simulated fluxes across different experimental conditions between day 2 and day 4. (Green) **Gln**$^-$ 2x$10^7$ LgEVs/mL vs control (Orange) **Gln**$^+$ 4x$10^7$ LgEVs/mL vs controls conditions.

The percentage difference columns highlight variations between MFI-predicted of LgEVs treated vs the respective control, noting instances of low flux or directional sign switches.

| | | Experimental Fluxes (nmol/E6 cells/day) | | | | MFA Calculated Fluxes (nmol/E6 cells/day) | | | | % Difference between +EV and control | | | |
|---|---|---|---|---|---|---|---|---|---|---|---|---|---|
| | | Control - Gln | -Gln +EV(2E7) | Control + Gln | +Gln +EV(4E7) | Control - Gln | -Gln +EV(2E7) | Control + Gln | +Gln +EV(4E7) | Control -Gln | -Gln +EV(2E7) | Control +Gln | +Gln +EV(4E7) |
| v0: | ATP + GLC --> G6P | | | | | 686 | 1544 | 993 | 1035 | | 124.9 | | 4.3 |
| v1: | G6P --> 3.0 ATP + 2.0 NADH + 2.0 PYR | | | | | 575 | 730 | 892 | 439 | | 26.9 | | -50.8 |
| v2: | NADH + PYR <=> LAC | | | | | 427 | 1596 | 371 | 639 | | 273.7 | | 72.4 |
| v3: | GLU + PYR <=> AKG + ALA | | | | | 70 | 67 | 426 | 643 | | -5.1 | | 50.8 |
| v4: | PYR --> AcCoA + CO2 + NADH | | | | | 1934 | 2105 | 1481 | 2069 | | 8.9 | | 39.7 |
| v5: | AcCoA + OXA --> CIT | | | | | 2295 | 2401 | 1966 | 2334 | | 4.6 | | 18.7 |
| v6: | CIT --> AKG + CO2 + NADH | | | | | 2295 | 2401 | 1966 | 2334 | | 4.6 | | 18.7 |
| v7: | AKG --> CO2 + NADH + SucCoA | | | | | 2314 | 2568 | 2300 | 2899 | | 11.0 | | 26.1 |
| v8: | SucCoA <=> ATP + SUC | | | | | 2349 | 2554 | 2341 | 2927 | | 8.7 | | 25.0 |
| v9: | SUC <=> FADH2 + FUM | | | | | 1995 | 2675 | 1849 | 3075 | | 34.0 | | 66.3 |
| v10 | FUM --> CO2 + PYR | | | | | 0 | 679 | 63 | 1091 | | | | 1626.4 |
| v11 | FUM <=> NADH + OXA | | | | | 2035 | 3158 | 1822 | 3975 | | 55.2 | | 118.2 |
| v12 | TRP --> ALA + 2.0 AcCoA + 4.0 CO2 + 2.0 NADH | | | | | 2 | 9 | 4 | 8 | | Very small flux | | |
| v13 | ASN <=> ASP + NH3 | | | | | 283 | 334 | 177 | 271 | | 18.2 | | 52.7 |
| v14 | AKG + ASP <=> GLU + OXA | | | | | 261 | -757 | 144 | -1641 | | Sign switched | | |
| v15 | NADH + PHE --> TYR | | | | | 0 | 23 | 0 | 28 | | Small flux | | |
| v16 | AKG + TYR --> Acetoac + CO2 + FUM + GLU | | | | | 14 | 52 | 13 | 58 | | Very small flux | | |
| v17 | CO2 + NADH + NH3 + SER <=> 2.0 GLY | | | | | -79 | 315 | -171 | 131 | | Sign switched | | |
| v18 | AKG + ATP + LEU --> AcCoA + Acetoac + FADH2 + GLU + NADH | | | | | 50 | 66 | 39 | 45 | | 32.8 | | 15.7 |
| v19 | THR --> AcCoA + GLY + NADH | | | | | 436 | 0 | 565 | 0 | | Very small flux | | |
| v20 | HIS --> GLU + NH3 | | | | | 4 | 0 | 0 | 0 | | Very small flux | | |
| v21 | GLN <=> GLU + NH3 | | | | | -101 | -42 | 244 | 421 | | -58.9 | | 72.3 |
| v22 | GLU <=> AKG + NADH + NH3 | | | | | 408 | -1890 | 234 | -2475 | | Sign switched | | |
| v23 | CYS --> NH3 + PYR | | | | | 864 | 10 | 0 | 24 | | Very small flux | | |
| v24 | AKG + ATP + VAL --> 2.0 CO2 + FADH2 + GLU + 3.0 NADH + SucCoA | | | | | 29 | 35 | 34 | 54 | | 19.5 | | 58.8 |
| v25 | AKG + ATP + ILE --> AcCoA + FADH2 + GLU + 2.0 NADH + SucCoA | | | | | 54 | 49 | 39 | 44 | | -9.5 | | 14.1 |
| v26 | THR <=> NADH + NH3 + SUC | | | | | -418 | 3 | -542 | 46 | | Sign switched | | |
| v27 | 2.0 ATP + MET + SER --> CYS + NADH + NH3 + SucCoA | | | | | 16 | 21 | 20 | 33 | | 26.9 | | 67.9 |
| v28 | Acetoac + SucCoA --> 2.0 AcCoA + SUC | | | | | 64 | 118 | 51 | 103 | | 83.3 | | 101.3 |
| v29 | 2.0 AKG + LYS --> 2.0 AcCoA + FADH2 + 2.0 GLU + 3.0 NADH | | | | | 17 | 30 | 22 | 47 | | 77.1 | | 115.1 |
| v30 | NADH + 0.5 O2 --> 2.5 ATP | | | | | 9639 | 9670 | 9281 | 10359 | | 0.3 | | 11.6 |
| v31 | FADH2 + 0.5 O2 --> 1.5 ATP | | | | | 2144 | 2854 | 1982 | 3265 | | 33.1 | | 64.7 |
| v32 | 0.01259 ALA + 0.01025 ARG + 0.00705 ASN + 0.01509 ASP + 0.6138 ATP + 0.05304 AcCoA + 0.002215 CHO + 0.00312 CYS + 0.01716 G6P + 0.01505 GLN + 0.00457 GLU + 0.01645 GLY + 0.00308 HIS + 0.00603 ILE + 0.01368 LEU + 0.01154 LYS + 0.00308 MET + 0.07733 NADH + 0.0025 NH3 + 0.00851 PHE + 0.01123 SER + 0.00855 THR + 0.00223 TRP + 0.00467 TYR + 0.00834 VAL --> Biomass + 0.00812 CO2 + 0.0039 FUM | | | | | 6483 | 2460 | 5869 | 2646 | | -62.1 | | -54.9 |
| v33 | 0.0096 ALA + 0.00835 ARG + 0.008 ASN + 0.00865 ASP + 0.8163 ATP + 0.0056 CYS + 0.009585 GLN + 0.0105 GLU + 0.01422 GLY + 0.0037 HIS + 0.00526 ILE + 0.01484 LEU + 0.012677 LYS + 0.00216 MET + 0.0068 PHE + 0.0232 SER + 0.01515 THR + 0.00402 TRP + 0.008658 TYR + 0.01824 VAL --> IgG | | | | | 354 | 469 | 219 | 168 | | 32.5 | | -23.3 |
| v34 | --> GLC | 683 | 2573 | 996 | 1837 | 686 | 1544 | 993 | 1035 | | | | |
| v35 | <=> LAC | -428 | -1140 | -371 | -598 | -427 | -1596 | -371 | -639 | | | | |
| v36 | <=> ALA | 13 | -36 | -354 | -579 | 13 | -40 | -354 | -616 | | | | |
| v37 | --> TRP | 18 | 27 | 18 | 15 | 18 | 16 | 18 | 14 | | | | |
| v38 | <=> NH3 | | | | | -1536 | 1344 | -720 | 2380 | | | | |
| v39 | --> ASN | 331 | 592 | 221 | 299 | 331 | 355 | 220 | 291 | | | | |
| v40 | <=> ASP | 79 | 60 | 58 | 52 | 79 | 51 | 58 | 51 | | | | |
| v41 | --> PHE | 56 | 79 | 52 | 62 | 58 | 47 | 51 | 52 | | | | |
| v42 | --> TYR | 48 | 73 | 42 | 51 | 48 | 44 | 42 | 44 | | | | |
| v43 | <=> SER | 436 | 749 | 351 | 478 | 436 | 449 | 350 | 455 | | | | |
| v44 | <=> GLY | -167 | -416 | -124 | -213 | -167 | -583 | -124 | -216 | | | | |
| v45 | --> MET | 37 | 49 | 38 | 44 | 37 | 29 | 38 | 41 | | | | |
| v46 | --> THR | 79 | 52 | 77 | 79 | 79 | 31 | 76 | 71 | | | | |
| v47 | --> ILE | 94 | 110 | 76 | 75 | 95 | 66 | 75 | 61 | | | | |
| v48 | --> VAL | 89 | 106 | 87 | 97 | 90 | 64 | 87 | 79 | | | | |
| v49 | --> CYS | | | | | 870 | 0 | 0 | 0 | | | | |
| v50 | <=> GLU | 130 | 105 | 104 | 109 | 130 | 63 | 104 | 107 | | | | |
| v51 | --> ARG | 88 | 134 | 77 | 97 | 88 | 80 | 77 | 80 | | | | |
| v52 | <=> GLN | 0 | 0 | 335 | 465 | 0 | 0 | 335 | 462 | | | | |
| v53 | --> HIS | 26 | 1 | 18 | 7 | 26 | 2 | 19 | 9 | | | | |
| v54 | --> LEU | 143 | 178 | 123 | 124 | 144 | 107 | 122 | 83 | | | | |
| v55 | --> LYS | 95 | 107 | 92 | 107 | 96 | 64 | 92 | 80 | | | | |
| v56 | --> O2 | | | | | 5892 | 6262 | 5632 | 6812 | | | | |
| v57 | <=> CO2 | | | | | -6755 | -6512 | -6124 | -6559 | | | | |
| v58 | <=> Biomass | 8104 | 4100 | 7336 | 6614 | -6483 | -2460 | -5869 | -2646 | | | | |
| v59 | <=> IgG | 443 | 782 | 274 | 421 | -354 | -469 | -219 | -168 | | | | |
| v60 | SER --> NH3 + PYR | | | | | 417 | 1619 | 431 | 1360 | | | | |
| v61 | ARG --> ORN + UREA | | | | | 19 | 1152 | 15 | 1973 | | | | |
| v62 | 2.0 ATP + CO2 + NH3 + ORN --> CLN | | | | | 0 | 1101 | 0 | 1922 | | | | |
| v63 | ASP + ATP + CLN --> ARG + FUM | | | | | 0 | 1101 | 0 | 1922 | | | | |
| v64 | AKG + ORN <=> 2.0 GLU + NADH | | | | | 19 | 51 | 15 | 51 | | | | |
| v65 | ATP --> | | | | | 26269 | 26269 | 26269 | 26269 | | | | |
| v66 | UREA <=> | | | | | 19 | 1152 | 15 | 1973 | | | | |
| v67 | G6P + 2.0 GLU --> 2.0 AKG + 2.0 NADH + 2.0 SER | | | | | 0 | 772 | 0 | 551 | | | | |
| | Total FADH2 flux produced | | | | | 2144 | 2854 | 1982 | 2566 | | 33.0691982 | | 29.45975914 |
| | Total NADH flux produced | | | | | 10470 | 15477 | 10337 | 10594 | | 47.82497016 | | 2.48946068 |
